# Fluke-borne viruses are a risk factor and diagnostic target for diseases caused by carcinogenic trematodes

**DOI:** 10.1101/2025.10.22.683923

**Authors:** Sujittra Chaiyadet, Tom Roblin, Anais Fauchois, Sarah Temmam, David Hing, Elise Jacquemet, Thomas Bigot, Blaise Li, Julia Kende, Lisandru Capai, Javier Sotillo, Thewarach Laha, Sophie Goyard, David Hardy, Thierry Rose, Alex Loukas, Paul J Brindley, Banchob Sripa, Nolwenn M Dheilly

## Abstract

Three trematode parasites, *Schistosoma haematobium*, *Opisthorchis viverrini,* and *Clonorchis sinensis* are recognized group 1 carcinogens for their roles in squamous cell carcinoma of the urinary bladder and bile duct cancer (cholangiocarcinoma). We aimed to assess the presence of fluke viruses and their association with malignancy in fluke infections. We showed that all three carcinogenic trematodes harbored related viruses. Focusing on *O. viverrini*, we observed virus persistence at discrete life stages and viral tropism for the fluke tegument, and we provided molecular and serological evidence of parasitized hosts’ exposure to viral RNA and proteins. Serological analyses confirmed seroconversion of residents in endemic sites and revealed that antibody titers and avidity against fluke-borne viruses increase in individuals with hepatobiliary diseases. Random forest models discriminated efficiently between liver-fluke-infected participants without disease and those with periductal fibrosis or cholangiocarcinoma. Thus, we show that fluke-borne viruses are risk factors and diagnostic targets for fluke-associated cancer.

## INTRODUCTION

Infectious diseases underpin more than 20% of cancers in the developing world ^1^. Although most cancer-causing pathogens are viruses, infection with several helminth parasites can also lead to cancer ^1–3^. The International Agency for Research on Cancer (IARC) has classified three eukaryotic parasites as group 1 carcinogenic agents: the liver flukes *Opisthorchis viverrini* and *Clonorchis sinensis*, which cause cholangiocarcinoma (CCA; a bile duct cancer), and the blood fluke *Schistosoma haematobium*, which causes squamous cell carcinoma of the urinary bladder (SCC). Both CCA and SCC have a dismal prognosis ^2,4–9^. Liver flukes afflict more than 40 million people in East Asia and Eurasia ^4,6,10–12^ whereas urogenital schistosomiasis affects 112 million people in 54 countries throughout sub-Saharan Africa, the Middle East, the Arabian Peninsula, and autochthonous cases have been reported in France since 2013 ^13–15^. Liver fluke infection may persist for up to 25-30 years and often remains asymptomatic. A subset of infected individuals displays a pro-inflammatory disease phenotype, with increased serum levels of Interleukin-6 (IL-6) and may develop advanced periductal fibrosis (APF) of the biliary tract, along with an elevated risk of CCA ^4,5^. This pro-inflammatory profile and persistent fibrosis may endure after anti-helminthic treatment with praziquantel has cleared the fluke infection ^16–19^. Without treatment, the asymptomatic excretion of *S. haematobium* eggs in the urogenital tract can persist for 3 to 5 years, causing hematuria and chronic inflammation, which in turn increases the risk of SCC ^20–22^. To date, control and treatment of these infections are achieved through regularly treating at-risk populations with preventive chemotherapy together with broader public health measures to reduce morbidity, even though reinfection may occur. Repeated treatment during childhood and in older age groups reduces the risk of severe disease later in life ^23–26^.

Recent studies revealed that viruses are ubiquitous in flatworms (phylum Platyhelminthes) ^27–32^. Trematodes are preferentially infected by viruses belonging to the orders *Mononegavirales, Hareavirales,* and *Martellivirales*, and, to a lesser extent, by viruses of the orders *Jingchuvirales* and *Picornavirales* ^28,32^. Of note, rhabdoviruses of trematodes and cestodes are excreted by these parasites, have frequently shifted host over the course of evolution, can infect helminth-infected vertebrate hosts, and occupy an ancestral position relative to most other viruses within the *Rhabdoviridae* ^28,32^. Together, these results indicate that viruses of parasitic flatworms pose a risk of infection to human hosts and may contribute to disease. However, the impact of fluke-borne viruses on parasite-associated diseases, including malignancy, remains unknown. Here, we show that all three carcinogenic flukes harbor viruses. Moreover, we demonstrate that opisthorchiasis triggers a robust immune response against *O. viverrini*-associated viruses that may contribute to the spectrum of infection-associated diseases, including APF and CCA.

## RESULTS

### The virome of carcinogenic flukes

The genomes of three viruses of *O. viverrini* and four viruses of *S. haematobium* were discovered through metatranscriptomics. Sanger sequencing and AmpliSeq complete genome sequencing were employed to complete and validate the genome composition of viruses of *O. viverrini* (Tables S1-S4). Detailed information on the viral genomes is provided in Table S5. Annotated viral genomes were compared to the previously reported genomes of viruses of *C. sinensis* ^28^ (Fig. 1). Phylogenetic analyses using the conserved RNA-dependent RNA polymerase (RdRp) confirmed that all viruses reported here are very closely related to other viruses previously known from trematodes, consistent with host specificity ^28,33^ (Fig. 1). More specifically, viruses of *O. viverrini* are closely related to viruses of *C. sinensis,* whereas viruses of *S. haematobium* are closely related to viruses discovered in other schistosome species| or in the small liver fluke *Dicrocoelium lanceatum* ^33^. Of interest, these viruses often show relatedness to viral taxa that include zoonotic pathogens highly pathogenic to humans, including *Togaviridae* (e.g., Chikungunya virus, Venezuelan equine encephalitis virus), *Alpharhabdoviridae* (e.g., Rabies virus, Mokola virus), and *Phenuiviridae* (e.g., Rift Valley fever virus, Heartland virus) (Fig. 1).

**Figure 1.**
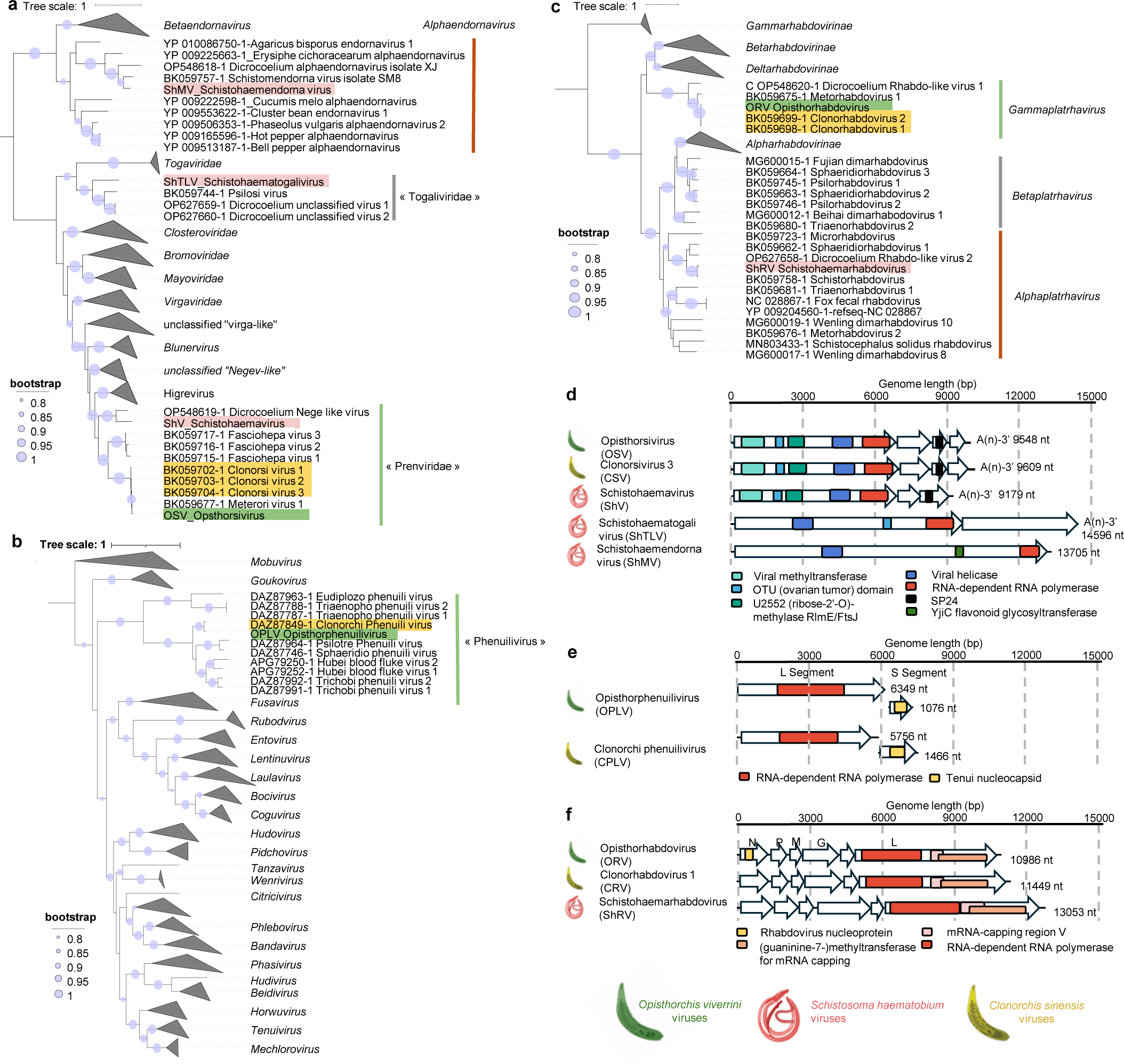
Discovery of viruses associated with carcinogenic parasitic trematodes. **a,c,e** Phylogenetic analyses of fluke viruses based on the RNA-directed RNA polymerases position them within the order *Martellivirales* (**a**), the family *Phenuiviridae* (**b**), and the family *Rhabdoviridae* (**c**). Viruses of *O. viverrini*, *C. sinensis,* and *S. haematobium* are highlighted in green, yellow and red, respectively. Phylogenetic trees were inferred after MAFFT alignment of the RdRp of RNA viruses with all viruses recognized by the International Committee for the Taxonomy of Viruses (ICTV) and additional unclassified viruses that are more closely related to the newly discovered viruses. Proposed names of novel families and genera reported in ^28^ are indicated in “brackets”. The trees were generated using PhyML with the LG substitution model. Branch points indicate the bootstrap results of Shimodaira-Hasegawa branch test with values >0.8. Genome composition of complete sequences of fluke viruses within the order *Martellivirales* (**d**), the family *Phenuiviridae* (**e**) the family *Rhabdoviridae* (**f**) and are depicted. Open reading frames were predicted using ORF finder and Conserved domains were predicted using NCBI conserved domain search.

Three viruses of *S. haematobium*, one virus of *O. viverrini* and one virus of *C. sinensis* belong to the order *Martellivirales* (Fig. 1a). The tentative species Opisthorsivirus (OSV), Clonorsivirus (CSV that comprises strains CSV-1, CSV-2, and CSV-3) and Schistohaemavirus (ShV) belong to a novel proposed family, named “Prenviridae” ^28^ that encloses only viruses of trematodes. The genome composition of prenviruses (Fig. 1d) is similar to mosquito-infecting negeviruses, including conserved domains for *Alphavirus* methyltransferase (pfam01660), the U2552 (ribose-2’-O)-methylase RlmE/FtsJ domain (pfam01728), the viral helicase (pfam01443), and RdRp (cd23254 corresponding to *Kitaviridae* RdRp) in ORF1, a putative glycoprotein composed of a signal peptide, transmembrane domain, and putative glycosylation sites in ORF2, and the putative virion membrane protein SP24 (pfam16504) in ORF3 (Fig. 1d). An additional ORF4 encoding a protein of unknown function is found in both OSV and CSV, but it is absent from ShV and from viruses of Fascioliidae . The Schistohaema togalivirus (ShTLV) belongs to a distinct clade named “Togaliviridae” of parasite-associated viruses that constitute to date the closest relatives of the zoonotic mosquito-borne *Togaviridae* family (Fig. 1a). The genome encodes conserved domains such as the RdRp (pfam00978) and the viral helicase (pfam01443) in the non-structural polyprotein, while the structural ORF encodes proteins of unknown function (Fig. 1d). The Schistohaemendornavirus (ShMV) falls within the genus *Alphaendornavirus* of vertically transmitted viruses (family *Endornaviridae*)(Fig. 1a). ShMV encodes a polyprotein harboring several functional domains, such as the typical RdRp of *Endornaviridae* members (cd23255), the viral helicase (pfam01443) and a glycosyltransferase (COG1819) (Fig. 1d).

The liver flukes *O. viverrini* and *C. sinensis* harbor the Opisthorphenuilivirus (OPLV) and the Clonorchi phenuilivirus (CPLV), respectively, which belong to a novel genera tentatively named “*Phenuilivirus”* ^28^, due to its close phylogenetic position to the arthropod-borne vertebrate viruses of the family *Phenuiviridae* (order *Hareavirales)* (Fig. 1b). Typical phenuivirus genomes are negative-sense, tri-segmented, although a few exceptions have just two segments. All known flatworm-associated phenuiviruses, including OPLV and CPLV, have a bi-segmented genome with the small S segment encoding the nucleoprotein (pfam05733), and the large L segment encoding the RdRp (pfam04196) (Fig. 1e). We deployed several strategies to identify a putative M segment encoding a glycoprotein including (i) a search for contigs containing panhandle sequences, hypothesizing that the M-segment panhandles of OPLV and CPLV would be similar to those flanking their L and S segments, and (ii) a search using relaxed similarities with known glycoproteins and M-segment nucleotide sequences from closely related species. All search strategies were unsuccessful, suggesting the absence of a third genome segment.

Finally, all three carcinogenic flukes host viruses that belong to the family *Rhabdoviridae*, order *Mononegavirales* (Fig. 1c,f). The Opisthorhabdovirus (ORV) and Clonorhabdovirus (CRV) belong to a novel genus of viruses of trematodes named *Gammaplatrhavirus* ^28^, whereas the Schistohaemarhabdovirus (ShRV) belongs to a separate genus of rhabdoviruses of neodermertan Platyhelminths named *Alphaplatrhavirus* ^28^. All three viral genomes encode the five canonical protein genes shared by all rhabodviruses: nucleoprotein N, phosphoprotein P, matrix protein M, glycoprotein G and RNA-dependent RNA polymerase (L), the latter harboring the typical functional domains of the *Mononegavirales* such as the RdRp (pfam00946) and the Guaninine-7-methyltransferase for mRNA capping (TIGR04198) (Fig. 1f). An additional accessory gene of unknown function positioned between the G and L genes encodes a putative viroporin ^34^. Of note, the position of this accessory gene is shared with other vertebrate-infecting viruses of the genera *Tibrovirus*, *Ephemerovirus*, and *Novirhabdovirus*, and the conserved ψ pseudogene remnant of an ancestral gene in *Lyssavirus* including rabies virus ^35–37^.

### Prevalence, quantification, and localization of *O. viverrini* viruses

RT-qPCR was performed on *O. viverrini* extracts from cercariae, newly excysted juveniles (NEJ) and adults confirmed virus persistence throughout the parasite life cycle (Fig. 2, Table S6-7). The viruses exhibited variable prevalence in adult flukes with 86% (19/22) of individual flukes infected with Opisthorhabdovirus (ORV), 13% (3/22) infected with Opisthorphenuilivirus (OPLV) and 55% (12/22) infected with Opisthorsivirus (OSV) (Fig. 2). Viral load per adult fluke was estimated for 16 individuals infected with at least one virus, and for a pool of cercariae and three pools of NEJ. Strikingly, although ORV and OPLV appeared to have variable abundance in flukes, with some individuals displaying minimal to undetectable levels of viral RNA, OSV was either absent or present in high abundance – an outcome indicating that the prevalence of ORV and OPLV in individual flukes may be underestimated. Viral loads ranged up to 1E+06 viral RNA copies of ORV, up to 1E+05 viral RNA copies of OPLV, and between 1+04 and 1+08 viral RNA copies of OSV per individual fluke (Fig. 2d). Viral RNA abundance in NEJ (50 ng RNA per pool) showed similar trends (OSV>ORV>OPLV), whereas the pool of cercariae differed, with more than 1E+09 ORV viral RNA copies detected in 50 ng of RNA input.

**Figure 2.**
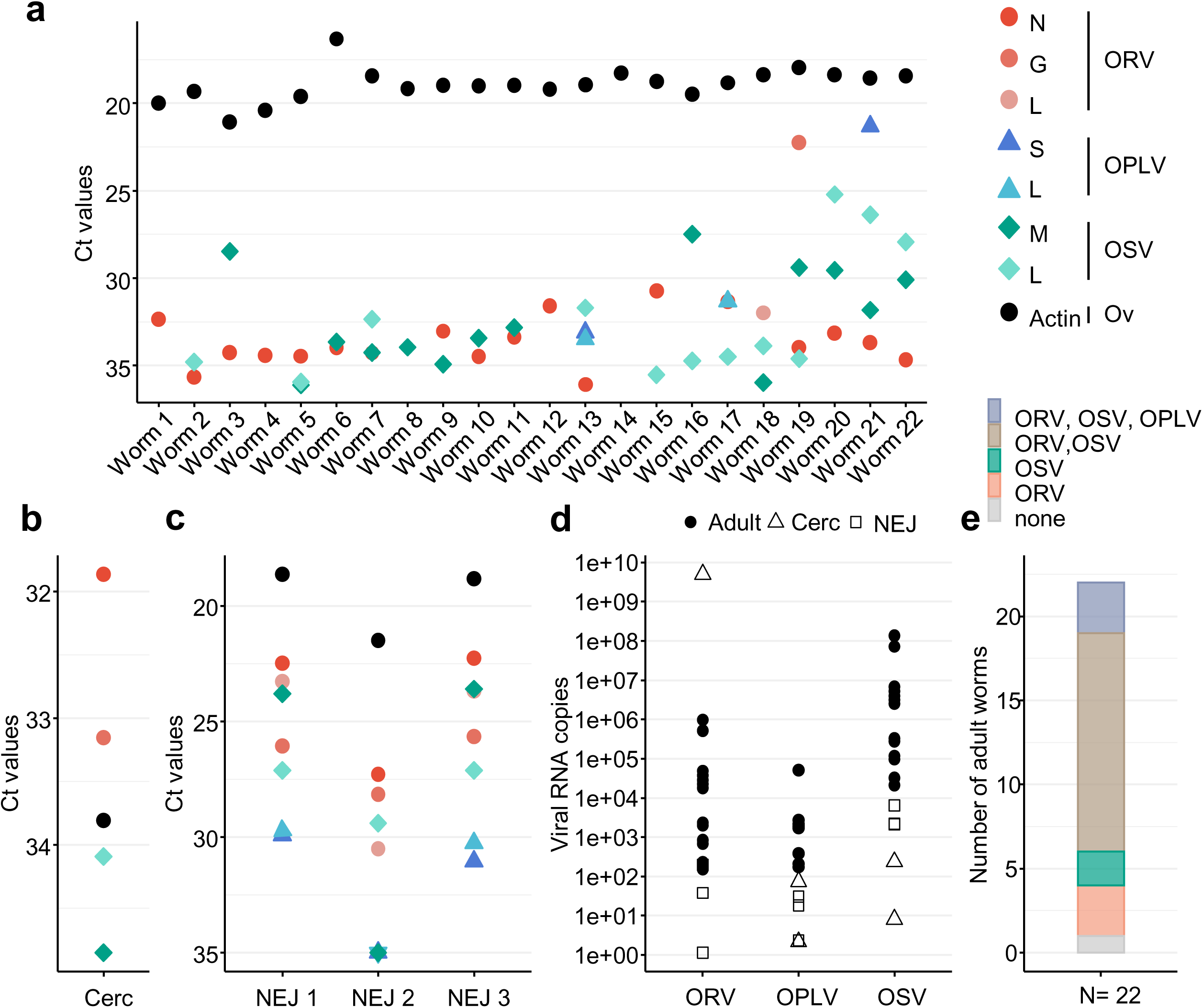
Molecular evidence supports the association of ORV, OSV and OPLV with *O. viverrini*. **a-c** RT-qPCRs were developed to detect RNA of ORV, OSV and OPLV in 22 individual adult flukes (**a**), cercariae (cerc; pool of 1000) (**b**) and newly excysted juveniles (NEJ; three pools of 250 each) with an equivalent of 50 ng of RNA input (**c**). **d** Viral RNA copies were estimated using a standard curve of synthetic RNA for 16 individual adult flukes (total RNA per individual, black circles), one pool of 1000 cercariae (50 ng of RNA, white square) and three pools of NEJ (50 ng of RNA, white diamond). We screened a total of 13, 9 and 13 individual worms for ORV, OPLV and OSV quantification, respectively. The median viral RNA abundance in individual adult worms was 1.7E+04 for ORV, 1.7E+03 for OPLV and 2.5E+06 for OSV. **e** The prevalence of virus infection in 22 individual adult flukes was calculated. Actin served as a positive control. Primers used for RT-qPCR and for synthetic RNA preparation are provided in Tables S6 and S7.

To determine viral tissue tropism in adult flukes, we used the branched-DNA fluorescence in situ hybridization approach that allows strand-specific nucleic acid detection ^38,39^ (Fig. 3-5, Table S8). The co-detection of both the positive- and negative-strand RNA enabled confident confirmation of probe specificity and indicated viral replication. Five individual adult flukes were examined, of which two were found negative for all three viruses, two were infected with all three viruses, and one was heavily infected with OSV. All three viruses were observed within *O. viverrini* eggs in the uterus of the parasite (Fig. 3-5). Both ORV and OSV were seen in cells of the outermost syncytial tegument of the flukes, and in the epithelia of both the gut and ovary (Fig. 4-5). ORV replication appeared notably elevated in the oral and ventral suckers (Fig. 4b and c), which is notable given the intimate association of the suckers of the adult worm with the epithelium of the parasitized bile duct and deeper periductal tissue. Although OSV was highly abundant in the tegument, the detection of (+)RNA throughout the fluke indicated virus presence in parenchymal cells embedding all other organs of adult flukes (Fig. 5).

**Figure 3.**
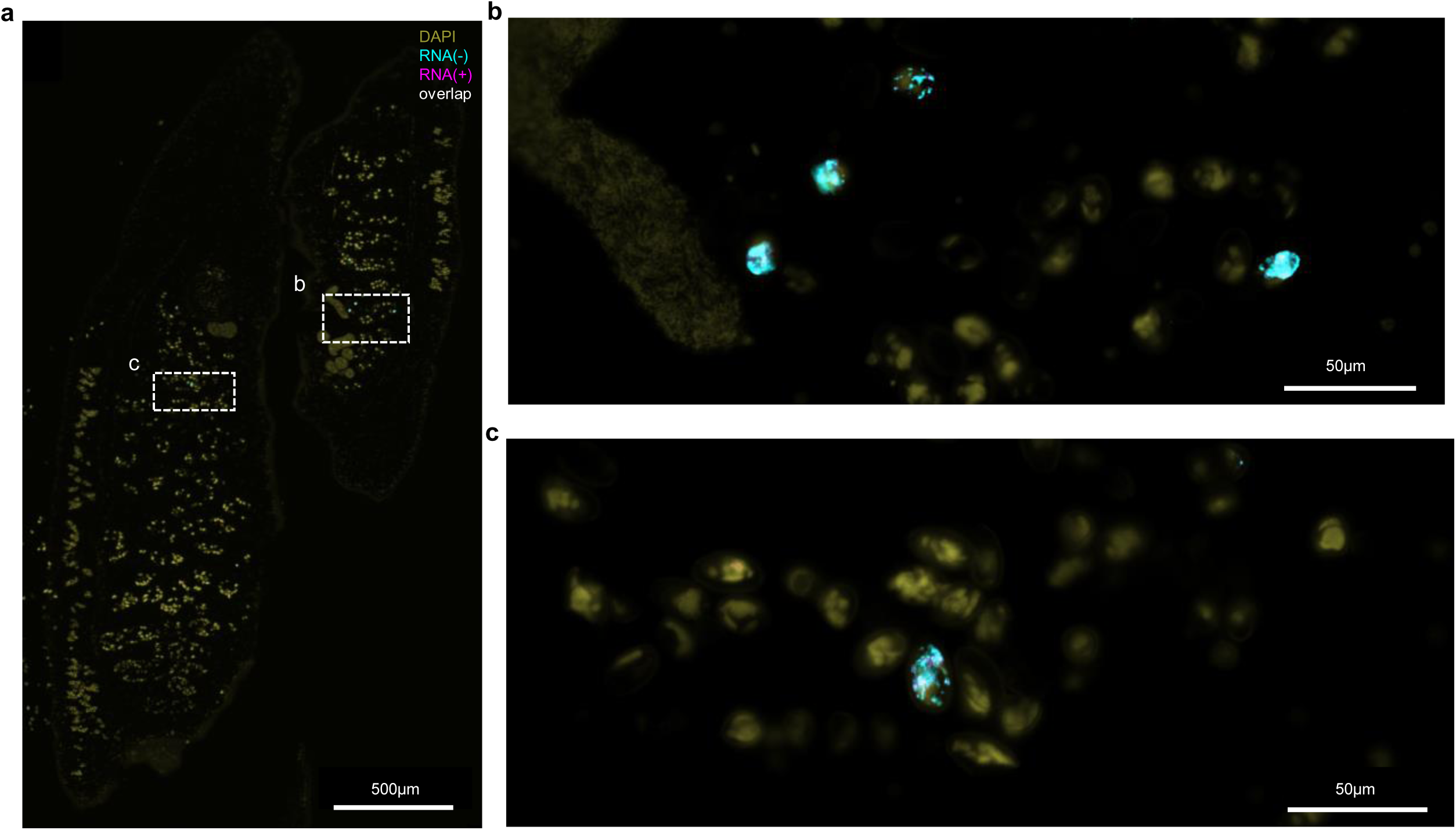
Localization of OPLV in *O. viverrini* by fluorescence immunochemistry. We used the RNAscope in-situ hybridization (ISH) assay to detect and visualize *O. viverrini* virus RNA in FFPE blocks of adult liver flukes. We used two set of ZZ probes binding to the positive-sense RNA, (+)RNA, in violet and to the negative-sense RNA, (-)RNA, in turquoise with co­localization observed in white (target sequences provided in Table S7). Nuclei were stained with DAPI and visualized in olive. OPLV is a negative-sense RNA virus, meaning the genome is (-)RNA; detection of (+)RNA indicates replication intermediates or viral transcription. **a** OPLV was observed in two individual worms **b-c** Co-localization of (+)RNA and (-)RNA revealed OPLV replication in fluke eggs (**a**) Scale bar: 500 µm. (**b-c**) Scale bar: 50µm.

**Figure 4.**
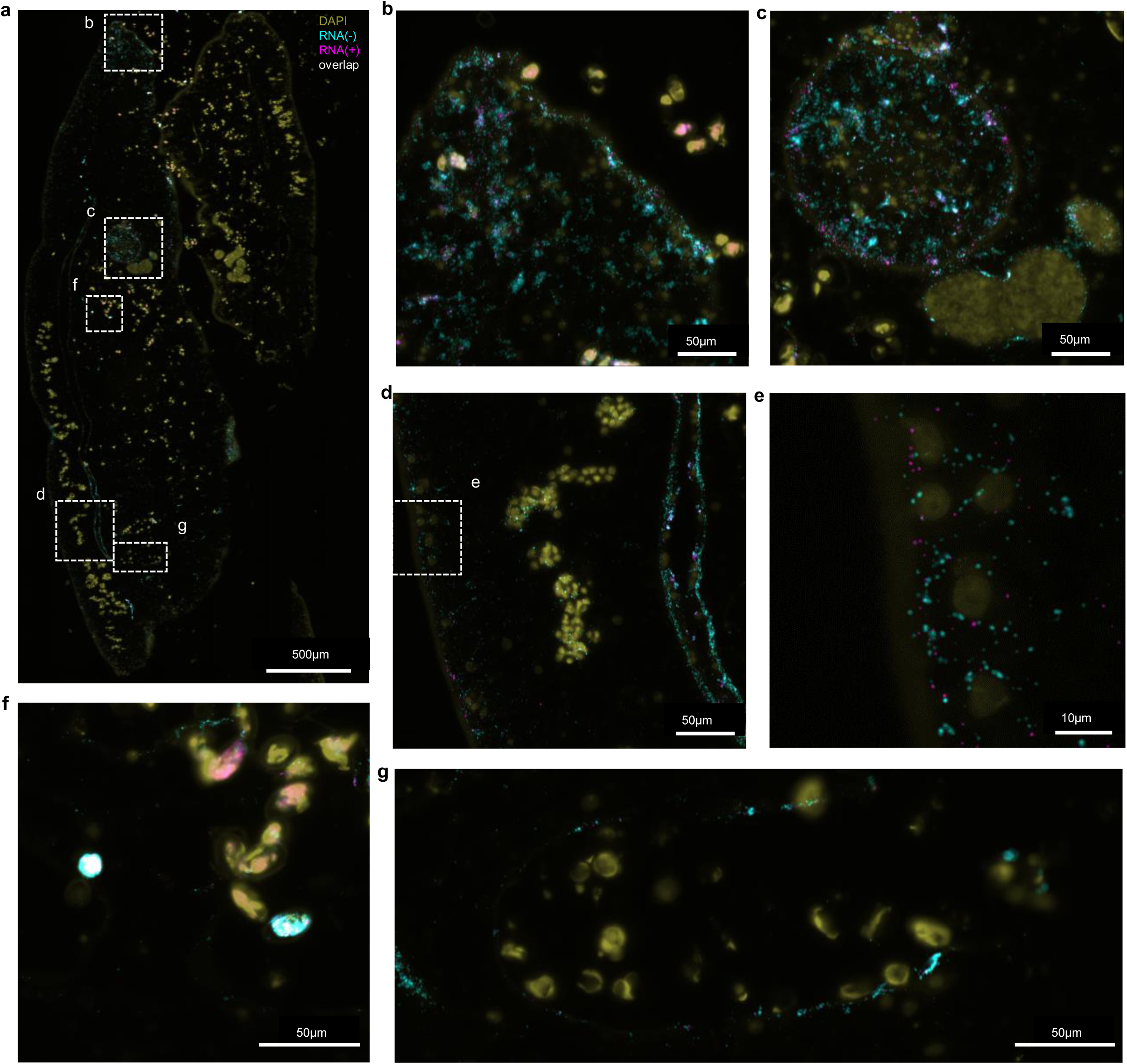
Localization of ORV in *O. viverrini* by fluorescence immunochemistry. We deployed RNAscope ISH to detect and visualize *O. viverrini* virus RNA in FFPE paraffin blocks of adult liver flukes. We used two set of ZZ probes binding to the positive-sense RNA, (+)RNA, in violet and to the negative-sense RNA, (-)RNA, in turquoise with co-localization observed in white (target sequences provided in Table S7). Nuclei were stained with DAPI and visualized in olive. ORV is a negative-sense RNA virus, meaning its genome is (-)RNA; detection of (+)RNA indicates replication intermediates or viral transcripts. **a** ORV was observed in two individual adult fluke with much greater virus abundance in the individual on the left. Co­localization of (+)RNA and (-)RNA revealed ORV replication in the oral sucker (**b**), ventral sucker (**c**), gut epithelium (**d**), tegument (**e**), eggs (f) and ovary epithelium (**g**). (**a**) Scale bar: 500 µm. (**b-d, f-g**) Scale bar: 50µm. (**e**) Scale bar: 10µm

**Figure 5.**
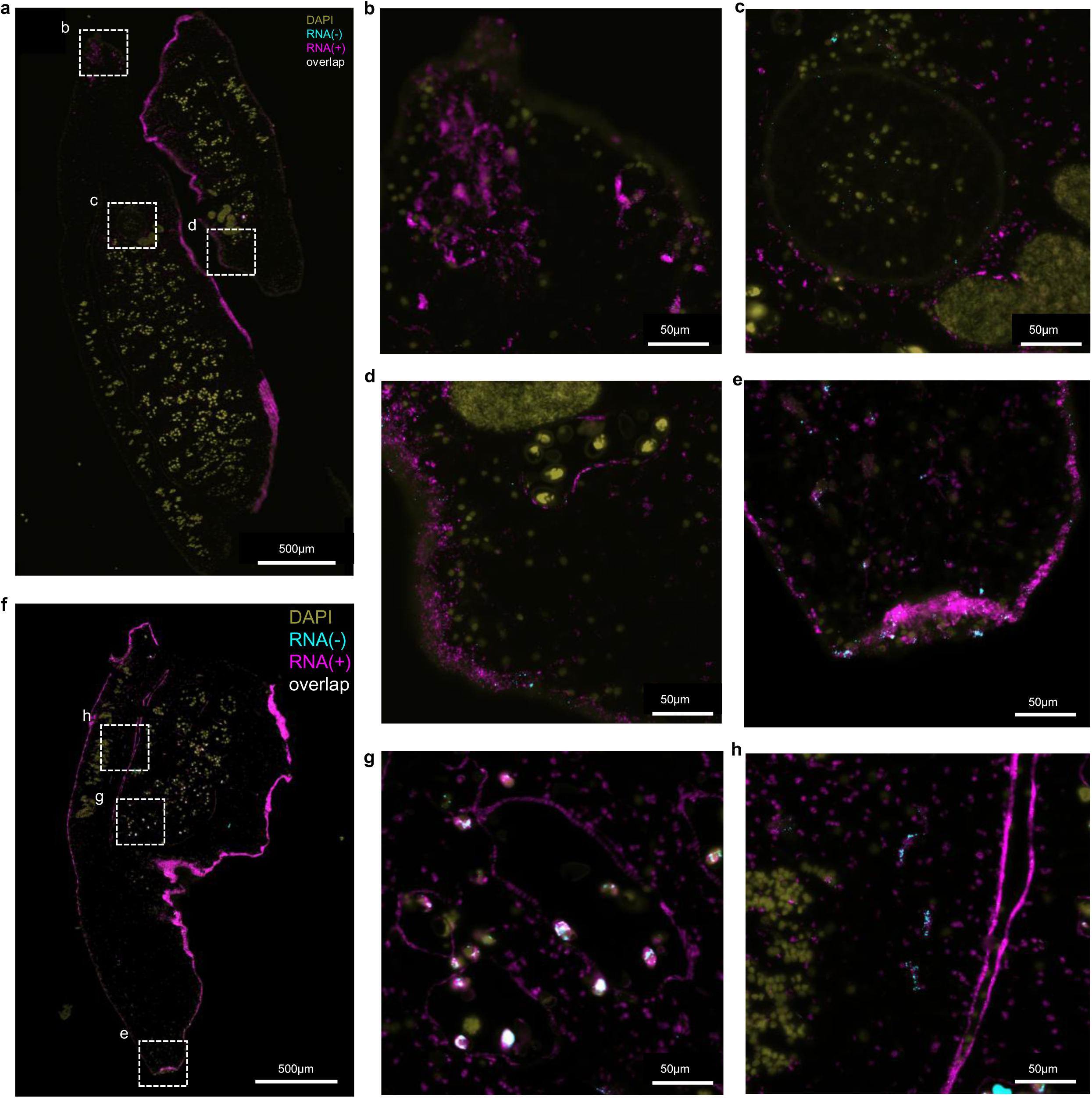
Localization of OSV in *O. viverrini* by fluorescence immunochemistry. We used the RNAscope in-situ hybridization (ISH) assay to detect and visualize *O. viverrini* virus RNA in paraffin blocks of adult liver flukes. We used two set of ZZ probes binding to the positive-sense RNA, (+)RNA, in violet and to the negative-sense RNA, (-)RNA, in turquoise with co­localization observed in white (target sequences provided in Table S7). Nuclei were stained with DAPI and visualized in olive. OSV is a positive-sense RNA virus, meaning its genome and transcripts are (+)RNA; detection of (-)RNA indicates replication intermediates. **a** OSV was observed in two individual adult flukes showing strong staining in the tegument. **f** a third individual displayed greater viral RNA abundance. Co-localization of (+)RNA and (-)RNA revealed OSV replication in the oral sucker (**b** and **e**), ventral sucker (**c**), tegument (**d**) eggs, ovary epithelium (**g**) and gut epithelium (**h**). Moreover, some cells with strong red signal indicating high replication foci were observed (**h**). (**a, f**) Scale bar: 500 µm. (**b-e, g-h**) Scale bar: 50µm.

### Extracellular vesicles mediate transfer of viral RNA

Two distinct types of extracellular vesicles (EVs) have been described in *O. viverrini* excretory/secretory (ES) products on the basis of their biogenesis and size: Large EVs (lEVs) are released from the plasma membrane by direct budding (diameter, 200 -1000 nm), whereas small EVs (sEVs) have an endocytic origin (40-150 nm)^40–42^. EVs are internalized by cholangiocytes and carry *O. viverrini* proteins and miRNA as cargo, which alter cholangiocyte physiology and phenotype ^40^. Total RNA and small RNA sequencing data from purified lEVs and sEVs reported in ^40^ were re-analyzed to assess the presence of viral nucleic acids. Total RNA sequencing data revealed that OSV was the most abundant virus in both sEVs and lEVs fractions, and a complete genome could be extracted from both (Fig 6a,b). In the sEV fraction, sequencing depth was homogeneous along the OSV genome, revealing the presence of complete viral genomic RNA (Fig. 6b). Traces of ORV and OPLV were also found in sEVs. However, it remains unclear whether OSV viral particles were co-purified with sEVs or whether the viral genome is also a component of sEV cargo.

**Figure 6.**
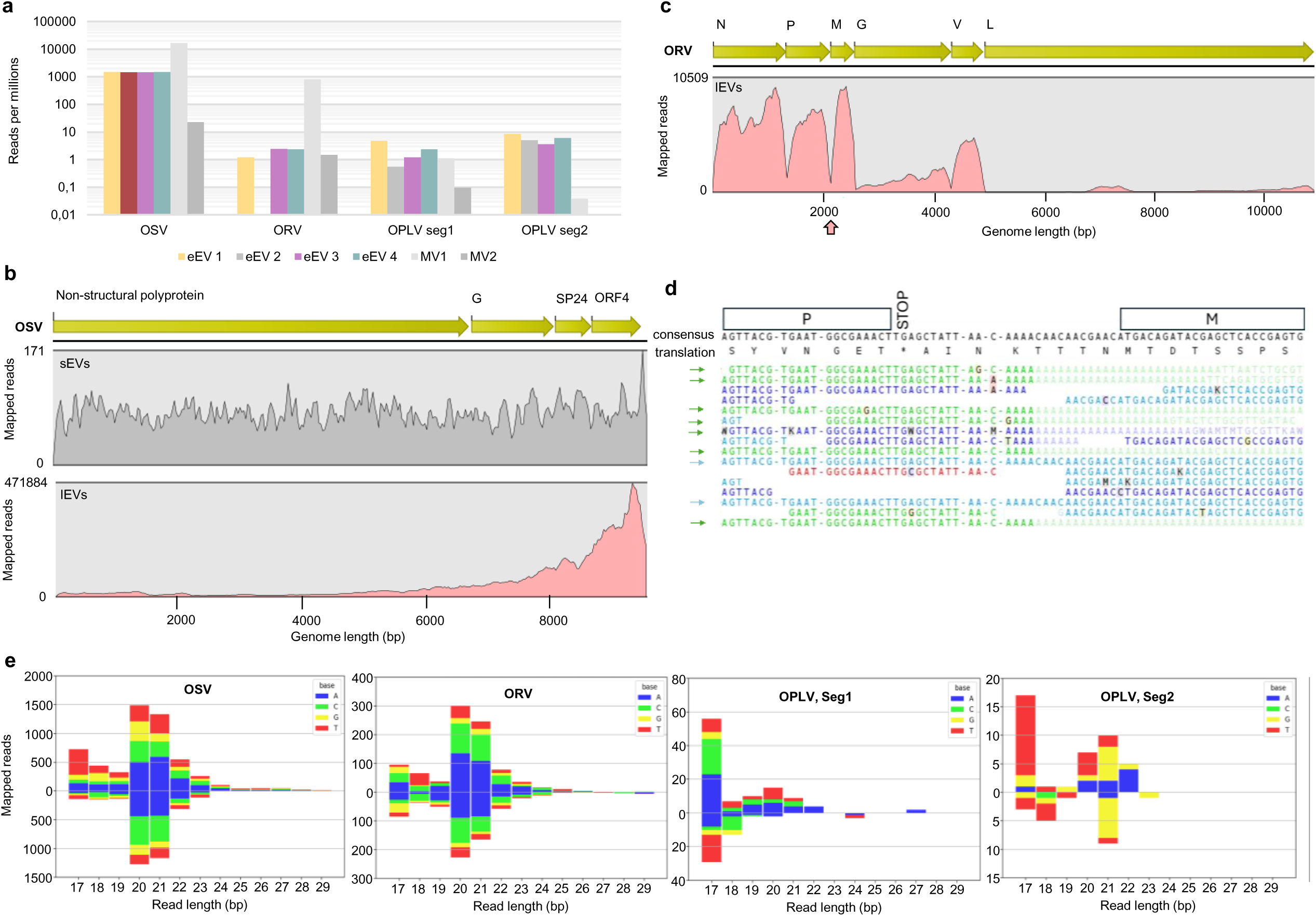
Extracellular vesicles mediate transfer of viral RNA. **a** Number of reads from small extracellular vesicles (sEV) and large EVs (lEV) *O. viverrini* samples mapped against the genomes of ORV, OSV and OPLV (Segment 1 and segment 2). Results are normalized in reads per millions of sequenced reads per sample. **b** Coverage of total RNA reads from sEVs and lEVs that mapped to the OSV genome with genome composition depicted. G: Glycoprotein. **c** Coverage of total RNA reads from lEVs that mapped to the ORV genome with genome composition depicted. N: nucleoprotein, P: phosphoprotein, M: matrix, G: glycoprotein, V: viroporin, L: large protein encoding the RdRp . **d** screenshot of the reads mapped to the intergenic region between the M and P genes of ORV. Green arrows point to mRNA reads with poly(A) tails whereas blue arrows point to genomic reads covering the intergenic region. **e** Size distribution, polarity and the 5’-terminal nucleotide of total viral small RNA from the sEV libraries. The plots show the cumulative values of two biological replicates. Plots obtained from each individual replicate library are provided as Fig. S1. The relative abundance of the different-size sense (top) and antisense (bottom) viral small RNAs is shown.

In lEVs, the OSV genome coverage was strongly biased towards the 3’ end, a result characteristic of subgenomic (+)RNA transcripts (Fig. 6b). A partial ORV genome (85% breadth of coverage) could also be obtained from lEVs, but coverage was uneven, with dips in intergenic regions (Fig. 6c). Focusing on these regions, we observed a small proportion of reads that mapped perfectly to the genome, confirming the presence of genomic RNA, and an abundance of reads with poly(A) tails characteristic of ORV mRNA (Fig. 6d). Together, these results indicate that ORV and OSV mRNA are packaged in lEVs. Again, it remains unclear whether ORV viral particles were co-purified with lEVs or whether the viral genome is also a component of lEV cargo. Of note, we also detected a partial ORV genome (75% breadth of coverage) and a complete OSV genome by mining previously reported mRNA sequence reads from *O. viverrini* (PRJEB65605) and observed the same mapping profiles characteristic of OSV subgenomic (+)RNA transcripts and ORV mRNA.

Viral small RNAs are typically produced by the antiviral defense system of invertebrates and display a size distribution of 22-23 nucleotides in nematodes and 21-22 nucleotides in arthropods ^43–45^. The role of the RNAi pathway role in antiviral defense has never been assessed in flatworms, including trematodes, but its viability was validated when siRNA was used to knock down gene expression in several species, including *O. viverrini* ^46–51^. Viral small RNAs mapping against all three viral genomes were detected in sEVs, but not in lEVs. We plotted the size distribution and 5’-terminal nucleotide composition of total 17 – 29 nt sense and antisense small RNAs mapped to ORV, OSV, and OPLV (Fig. 6e). There was a predominance of viral short interfering RNAs (vsiRNAs) of 20-21 nt, in the size range of Dicer products, mapping in equal abundance to the genomic and antigenomic strands of ORV and OSV, with a slight bias against G and, to a lesser extent, toward T at the 5’ position (Fig. 6e). These findings indicate that both ORV and OSV infections are active in adult *O. viverrini* worms and provide the first indication that small RNAs contribute to the antiviral defense system in the Trematoda. In contrast, for OPLV, a smaller proportion of 20-21 nt long vsiRNAs mapped against either Seg1 or Seg2 of the viral genome. This result is consistent with the observation that OPLV replicates within eggs rather than in the tegument.

To investigate transmission of fluke viruses to fluke-infected mammalian hosts, we deployed molecular approaches for which we collected tissues from five individual hamsters experimentally infected with *O. viverrini* one year prior. Using RT-qPCR, nucleic acids of all three viruses were detected in the liver, kidney, and stool pellets of these hamsters, and were confirmed through Sanger sequencing. Of note, however, we observed late Ct values (Ct > 35) in most samples, around or beyond the limit of detection of the assays, so that the presence of viral nucleic acids and virus transmission could not be definitively established (Fig. S2).

### Hamsters infected with *O. viverrini* develop IgG antibodies to fluke viruses

To investigate the exposure of fluke-infected hosts to fluke viral proteins, we deployed serological approaches and collected sera from the five individual hamsters from day 1 to day 360 after experimental infection with *O. viverrini*. We investigated seroconversion during *O. viverrini* infection to fluke and viral antigens using a luciferase-linked immunosorbent assay (LuLISA) ^52^. We observed high titers of IgG antibodies to the fluke antigen *Ov-*TSP-2 (tetraspanin) ^53^, the ORV nucleoprotein, the OPLV nucleocapsid, and OSV SP24 in the sera with peak values at about 100 days of infection with *O. viverrini* (Fig. 7a). The longitudinal profile demonstrates the transition from a baseline signal at day 1 to a sustained antibody response peaking at day 90-120 days later, thereby confirming seroconversion and the specificity of this LuLISA assay (Fig. 7a). The IgG titer at day 90 post-infection (Fig. 7b) was determined using an endpoint titration method, in which the positivity threshold for each hamster was defined as the highest serum dilution yielding a signal greater than the corresponding baseline on day 1 post-infection (Fig. S3). This approach revealed individual variability, with lower IgG titers against fluke viruses matched by lower IgG titers against the *O. viverrini* antigen in hamster H1.

**Figure 7.**
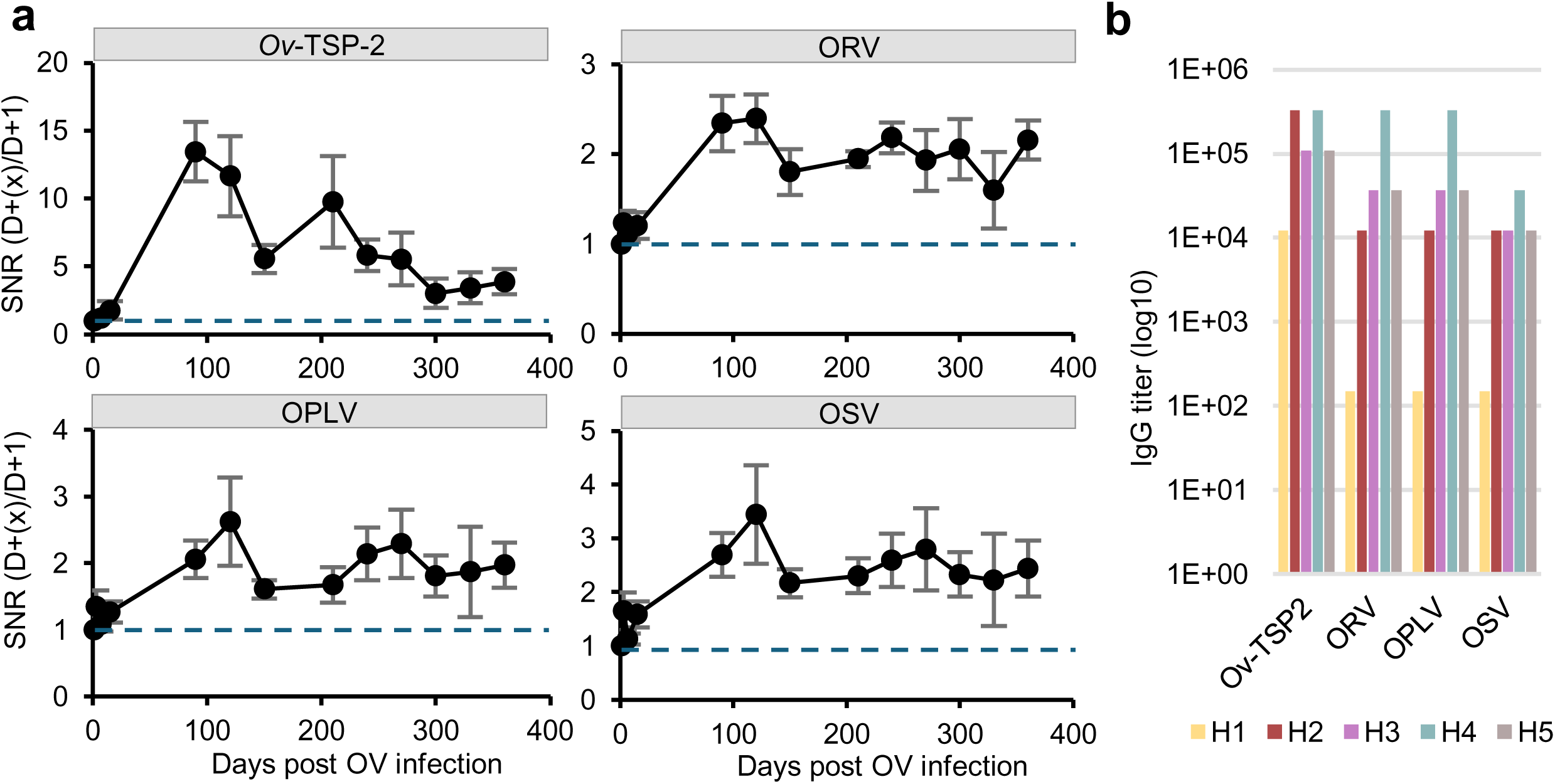
Serological detection of fluke viruses in parasitized hamsters. **a** longitudinal IgG response of *O. viverrini* infected hamsters over the 12 months study period, as the mean fold­change relative to the day 1 (D1) baseline ± SEM (n=5). **b** serum levels of anti-*Ov*-TSP-2, anti-ORV anti-OSV and anti-OPLV IgG in the hamsters. The reported antibody titer thus represents the final dilution factor at which the D90 serum remained above the signal at D1 post-infection.

### Seroconversion in *O. viverrini*-infected humans

Next, we investigated whether naturally acquired *O. viverrini* infection of humans was associated with the development of antibodies against ORV, OPLV, and OSV in the sera of 555 independent *O. viverrini*-infected and non-*O. viverrini*-infected control study participants. Sera from 455 *O. viverrini*-infection cases were retrieved from the biobank of the Tropical Disease Research Center (TDRC), Faculty of Medicine, Khon Kaen University, Thailand. Because *O. viverrini* is highly prevalent in the study region, and individuals may retain antibodies for prolonged periods after parasite clearance or praziquantel treatment, sera from 100 residents of France (here termed non-endemic French subjects) were used to set cut-off values. Antibody responses against Ov-TSP-2, ORV, OPLV, and OSV were significantly higher in *O. viverrini*-infected individuals from Thailand than in non-endemic French controls (Wilcoxon-Mann-Whitney test, all p < 0.0001), with moderate effect sizes (r = 0.43–0.49, 95% CI : 0.37–0.54; Fig. 8a, Table S9). To assess diagnostic performance, the results were further analyzed by receiver­operating characteristic (ROC) curve analysis (Fig. 8b, Table S10),yielding an area under the curve (AUC) of 0.86 (Sensitivity, Se 74%, Specificity, Sp 94%) for *Ov-*TSP-2, 0.83 (Se 74%, Sp 85%) for ORV, 0.87 (Se 71%, Sp 92%) for OSV, and 0.82 (Se 76%, Sp 82%) for OPLV (Table S10). Seroconversion was established using the threshold calculated from ROC curve analysis. Seroconversion against *Ov-*TSP-2, ORV, OPLV, and OSV were more prevalent in infected individuals (335/455, 75% for TSP-2; 346/455, 78% for ORV; 342/455, 77% for OPLV and 317/455; 71% for OSV) than in non-endemic controls (8, 21, 19 and 9 /100, respectively, Fig. 8a), confirming that *O. viverrini* infection was positively associated with ORV, OPLV, and OSV seroconversion.

**Figure 8.**
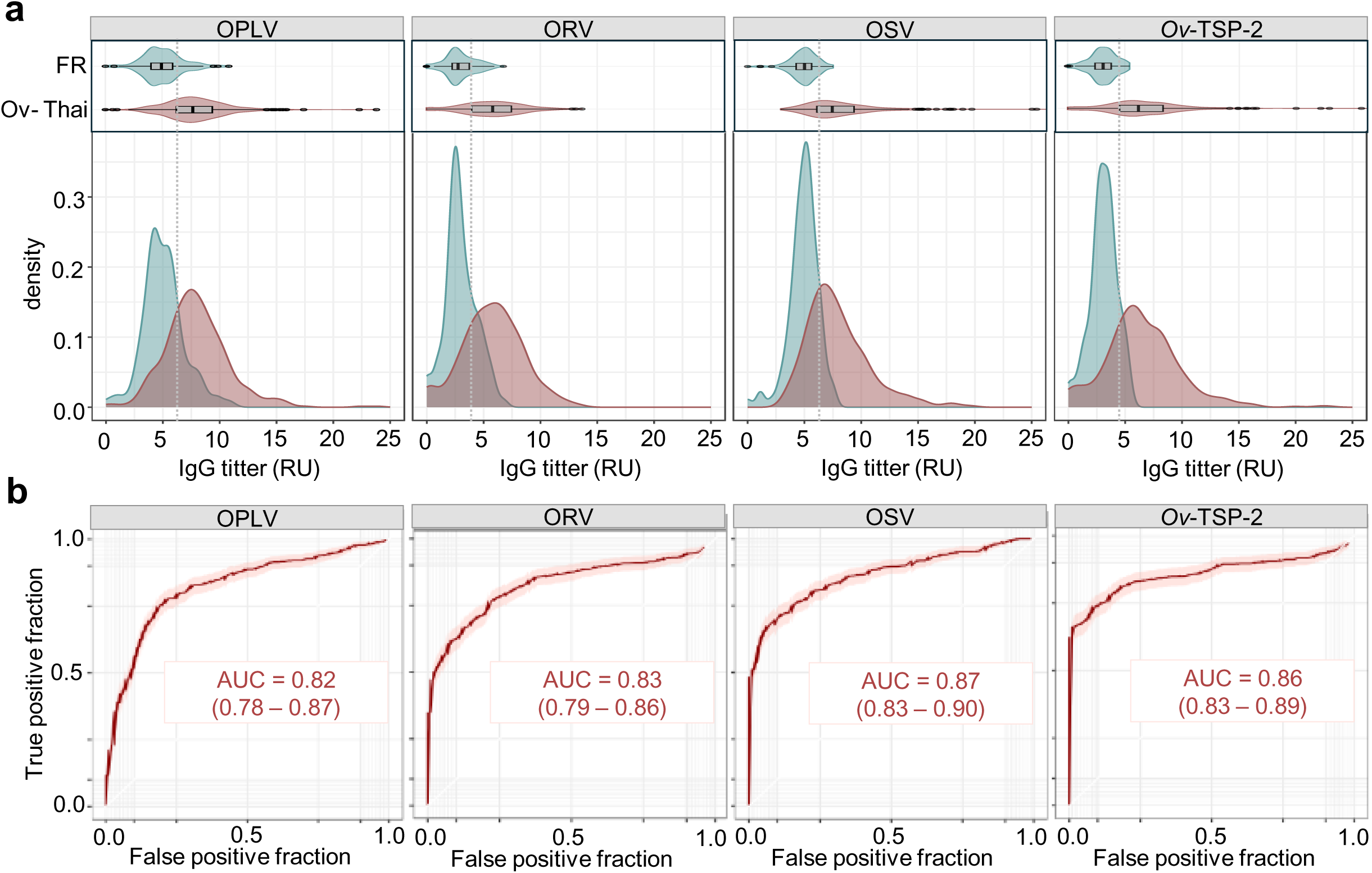
IgG titers for viral antigens increased in participants with past or current *O. viverrini* infection. **a** quantification of IgG titers against OPLV, ORV, OSV and *Ov-*TSP-2 in French non-endemic subjects (FR, *n*=100) and Thai *O. viverrini* infected subjects (Ov-Thai, n=455). Two-sided Wilcoxon-Mann-Whitney tests show significantly higher titers in Ov-Thai than in FR for all four antigens (all *p* < 0.001 after Benjamini-Hochberg correction), with moderate effect sizes (Table S9). The grey dashed line indicates the threshold calculated by receiver-operator characteristics (ROC) curve analysis. **b** ROC curve analysis displaying the Area Under the Curve (AUC) and confidence interval (Table S10).

### Elevated IgG titer against fluke viruses is associated with increased risk of advanced periductal fibrosis

As established in both in humans and rodent models, hepatic periductal fibrosis is a precursor of liver-fluke-induced CCA ^4,54,55^. Thus, we sought to determine if antibody titers against fluke viral proteins were elevated in *O. viverrini*-infected individuals with liver diseases. A case-control study was designed to investigate the association between serum antibodies and APF in serum samples collected from a community-based opisthorchiasis control program in Northeast Thailand ^56^. The study included 184 *O. viverrini*-positive cases without APF (APF-) and 184 *O. viverrini*-positive cases with APF (APF+). There were no significant differences in the demographic characteristics of participants in these two groups, including age, sex, and *O. viverrini* egg count (egg per gram (EPG) of feces). Antibody titers against ORV, OSV and OPLV were higher in sera of APF+ subjects than in those of APF-subjects (Wilcoxon-Mann-Whitney tests, *p*-value <0.001 for each comparison) with large effect size for ORV (r=0.5 95% CI: 0.42-0.58), moderate for OSV (r=0.42 95% CI: 0.33-0.51) and small for OSV (r=0.28 95% CI: 0.18-0.37)(Fig. 9a, Table S11, Table S12). In contrast, there was no difference in the antibody titers against *Ov-*TSP-2 between APF+ and APF-subjects (p=0.737, r=0.02 95% CI: 0­0.13), Fig. 9a, Table S12). To determine whether antibody responses were associated with APF, we performed multivariable logistic regression analysis, adjusting for age, sex, and *O. viverrini* EPG. Antibodies to ORV (adjusted OR = 2.05, 95% CI: 1.66–2.52), OPLV (OR = 1.70, 95% CI: 1.43–2.00), and OSV (OR = 1.19, 95% CI: 1.07–1.32) were significantly associated with increased odds of APF (all *p*< 0.001). Conversely, antibody levels to *Ov-*TSP-2 were negatively associated with APF (OR = 0.79, 95% CI: 0.71–0.89; *p*-value < 0.001).

**Figure 9.**
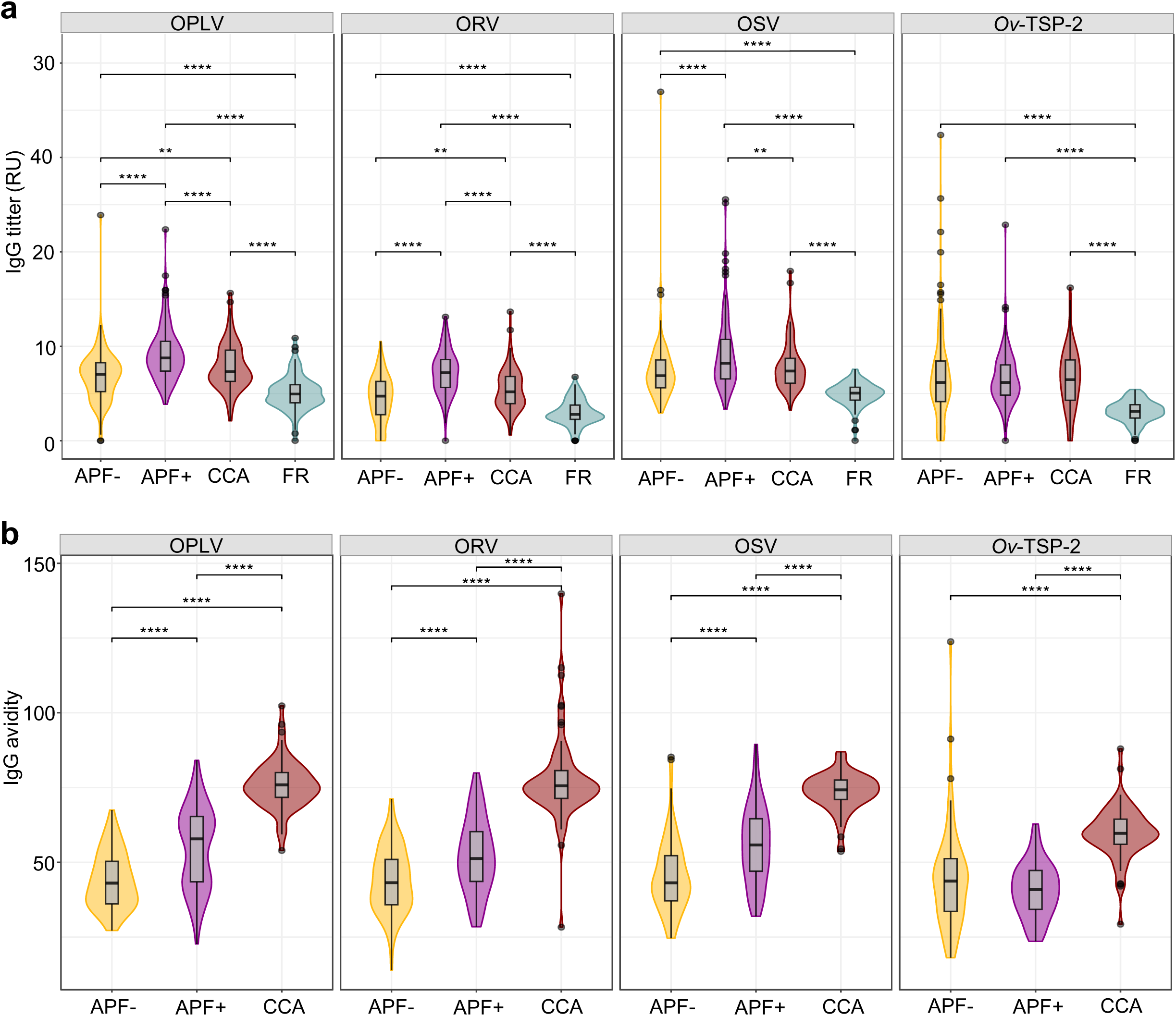
IgG titers and IgG avidity in non-endemic, and *O. viverrini* infected study participants with APF-, APF+ and CCA phenotypes. **a** IgG titers against OPLV, ORV, OSV and *Ov-*TSP-2 in non-endemic French subjects (FR, *n*=100) and Thai subjects without periductal fibrosis (APF-, *n*=184), with periductal fibrosis (APF+, *n*=184) and with cholangiocarcinoma (CCA, *n*=71). **b** IgG avidity index against OPLV, ORV, OSV and *Ov-*TSP-2 in Thai subjects without periductal fibrosis (APF-, n=72), with periductal fibrosis (APF+, *n*=72) and with CCA (*n*=71). Kruskal-Wallis test for differences among groups was significant (*p*-value<0.001)(Table S11-S12). Two-sided Wilcoxon-Mann-Whitney tests for difference between groups adjusted with BH correction (Table S13-S14) are displayed on the plot with the following code: “ ” if *p*-value>0.05, “*” if *p*-value<0.05, “**” if *p*-value<0.01, “***” if *p-*value<0.001, “****” if *p*-value<0.0001.

### Elevated IgG avidity against fluke viruses is associated with increased risk of cholangiocarcinoma

An additional 72 Thai patients with CCA were screened for serum antibody responses to viral and fluke antigens. Antibody titers against *Ov*-TSP-2 in sera of *O. viverrini*-infected cases did not differ among hepatobiliary disease states (APF-, APF+ and CCA) (Wilcoxon-Mann-Whitney tests, all p=0.737, r=0.02-0.03 95% CI: 0-0.16, Fig. 9a, Table S12). However, antibody titers were slightly higher in the CCA group than in the APF-group for ORV (*p*-value = 0.009, r=0.16 95% CI: 0.04-0.27) and OPLV (*p*-value =0.009, r=0.16 95% CI: 0.04-0.27) but not for OSV (*p*-value =0.061, r=0.11 (0.0065-0.23)) and were significantly lower in CCA subjects than in APF+ subjects for ORV (*p*-value <0.001, r=0.37 95% CI: 0.27-0.48), OPLV (*p*-value <0.001, r=0.24 95% CI: 0.11-0.35), and OSV (*p*-value =0.009, r=0.16 95% CI:0.05-0.28) (Fig. 9a, Table S12). This indicates that we could not differentiate among APF-, APF+ and CCA cases solely based on IgG titer.

The IgG avidity - the strength of binding between multivalent antigens and IgG antibodies - increases with maturation of the immune response and depends on the initial infectious dose or rate of reinfection^57^. The addition of urea in the incubation step is a simple but efficient means to reduce antibody binding and measure the avidity index for individual samples, and it has been proposed as a reliable method for diagnosis ^58–61^. The IgG avidity index was measured in sera of 71 CCA patients, and 72 subjects from each of the APF- and APF+ cohorts (Fig. 9b). APF+ individuals had IgG with higher avidity than APF-individuals for ORV (*p*-value <0.001, r=0.33 95% CI: 0.18-0.48), OPLV (*p*-value <0.0001, r=0.41 95% CI: 0.28-0.54) and OSV (*p*-value <0.0001, r=0.39 95% CI: 0.26-0.53), but not for *Ov*-TSP-2 (*p*-value=0.309, r=0.08 95% CI: 0­0.23) (Kruskall-Wallis & Wilcoxon-Mann-Whitney tests, Table S13, Table S14). The IgG avidity further increased in CCA patients and was largely higher than in APF+ cases for ORV (*p*-value <0.0001, r=0.75 95% CI: 0.66-0.81), OPLV (*p*-value <0.0001, r=0.72 95% CI: 0.63-0.8), OSV (*p*-value <0.0001, r=0.67 95% CI: 0.56-0.76), and *Ov*-TSP-2 (*p*-value <0.0001, r=0.75 95% CI: 0.67-0.82) (Fig 9b, Table S14).

### Establishing predictive models for advanced periductal fibrosis and cholangiocarcinoma on the basis of serology results

Next, we used random forest models, which add substantial value to diagnostic applications by integrating results from multiple serological assays into a single, unified prediction. We first tested the sensitivity and specificity of models that used IgG titers for all antigens together (ORV, OSV, OPLV, and *Ov-*TSP-2) to discriminate between non-infected and infected subjects. We used a randomly selected subset of 385 sera for training and 170 for validation of the models (Fig. 10 a-c, Table S15). The resulting model had an overall accuracy of 94% (89-97%) (significantly different from the null model, *p-*value = 0.0000055), which was superior to the accuracy of the test based on the fluke antigen *Ov*-TSP2. While the *Ov*-TSP2 IgG titer was the most important variable for the model, OSV, ORV, and OPLV IgG titers maintained > 20% importance in the model (Fig. 10c, Table S17). A second model that used only IgG titers for viral antigens only (ORV, OSV, and OPLV) yielded a slightly lower accuracy of 89% (84-94%) (significantly different from null model, *p*= 0.0075145) but remained more accurate than a test based on the fluke antigen *Ov*-TSP2 alone (Fig. 10, Table S15).

**Figure 10.**
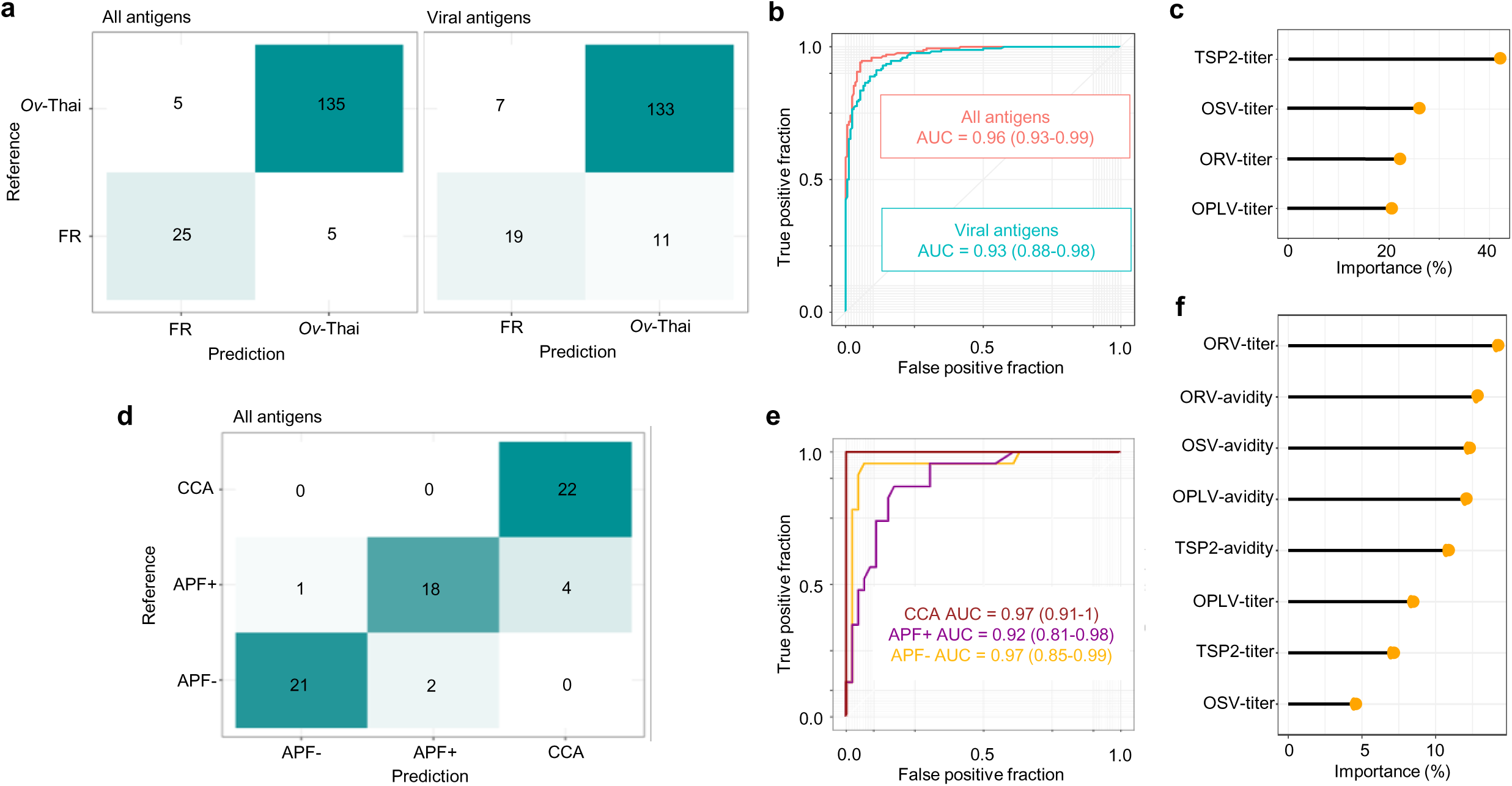
Random forest models based on IgG titers and IgG avidity discriminate among non-endemic, and *O. viverrini* infected subjects with APF-, APF+ and CCA phenotypes. **a-c** We used random forest models to classify French non-endemic controls and Thai *O. viverrini* parasitized participants based on IgG titers using a training set of 220 patients and a validation set of 110 patients. **a** The confusion models show prediction results on the validation set for a model using all antigens (ORV, OPLV, OSV and *Ov*-TSP2) and a model using viral antigens only (ORV, OSV and OPLV). **b** Results were further analyzed using ROC curve analysis. **c** displays the importance of the variables in the model using all antigens (Table S17). **d-f** We used a random forest model to classify APF-, APF+ and CCA subjects based on IgG titers and IgG avidity using a training set of 147 subjects and a validation set of 68 subjects. **d** The confusion model shows prediction results on the validation set. **e** Results were further analyzed using ROC curves. **f** displays the importance of the variables in the model (Table S17).

Finally, we tested the accuracy of a random forest model using both IgG titers and IgG avidity for all four antigens to differentiate among hepatobiliary disease states (APF-, APF+, and CCA) in *O. viverrini*-infected cases. We used a randomly selected training set of 147 individuals and a validation set of 68 individuals. The accuracy of the resulting model was 89.7% (79.9-95.8%) (significantly different from the null model*, p-*value = 0.000) (Table S16). Results were analyzed using ROC curves showing a prediction with high sensitivity (Se) and high specificity (Sp): an AUC of 1.0 for the detection of CCA (Se 100%, Sp 91%), an AUC of 0.89 for the detection of APF+ (Se 78%, Sp 96%), and an AUC of 0.95 for the detection of APF-(Se 91%, Sp 98%) (Fig. 10, Table S16). The paramount variables for the model were ORV IgG titer, followed by ORV, OSV, and OPLV avidity indices (Fig. 10e, Tables S17).

## DISCUSSION

Herein, we report the discovery of novel viruses associated with the food-borne Asian liver fluke, *Opisthorchis viverrini* and with the blood fluke *Schistosoma. haematobium*. Focusing on *O. viverrini*, we present laboratory-based evidence that parasitized individuals are exposed to both viral RNA and proteins and develop a corresponding humoral immune response. Using cohorts of *O. viverrini*-infected study participants, we established that infected individuals seroconverted against all three viruses and showed that infection by fluke viruses is a risk factor for disease. Indeed, our random forest models showed that serological assays targeting fluke viruses alongside *O. viverrini* distinguished between uninfected and infected subjects and, of potentially greater clinical importance, differentiate among patients according to hepatobiliary disease status.

Each of these three carcinogenic trematodes is associated with several viruses, including members related to virus families that contain zoonotic pathogens highly pathogenic to humans, such as alphaviruses (e.g., Chikungunya virus, Venezuelan equine encephalitis virus), alpharhabdoviruses (e.g., Rabies virus, Mokola virus), and phenuiviruses (e.g. Rift valley Fever virus, Heartland virus). All viruses discovered in *S. haematobium* and *O. viverrini* were closely related to other viruses previously discovered in other species of trematodes ^28,33^. Moreover, we showed that ORV and OSV are processed by the *O. viverrini* RNA interference pathway ^51,62^ confirming that these viral infections are active, but also under control by the fluke. This is the first report that this pathway is involved in antiviral defense by a trematode.

Virus tropism and transmission are key features that can help determine the zoonotic risk posed by novel parasite-associated viruses ^63^. All three viruses of *O. viverrini* were observed replicating in eggs, as assessed by the co-detection of both (-)RNA and (+)RNA, which suggests vertical transmission of the viruses. Moreover, the viruses persist throughout the developmental life stages of the fluke. In concordance, virus persistence at different life stages and vertical transmission were previously shown in *Schistocephalus solidus* and *Schistosoma japonicum* ^32,64^. Vertical transmission may prove to be a common feature of most viruses of Neodermata as an adaptation to parasitism that limits opportunities for horizontal transmission. Notably, however, the detection of both ORV and OSV in the tegument and EVs of *O. viverrini* suggests the potential for horizontal transmission of these viruses. We previously reported the presence of the rhabdovirus of *S. solidus* was found in several tissues of parasitized hosts, indicative of viral transmission ^32^. Moreover, rhabdoviruses of Neodermata do not display co-diversification with their hosts, which implies that host shifts are frequent during the evolution of these viruses ^28^. In contrast, OPLV was detected exclusively in eggs, which suggests strict vertical transmission. Moreover, neither OPLV nor any other related viruses detected in Neodermata possess a segment M encoding the glycoprotein, and all these Phenuiliviruses co-diversify strongly with their parasitic hosts ^28^. Finally, a recent report of viruses of the blood fluke *S. japonicum* in Hubei Province, China, further demonstrated that the viruses play a critical role in oogenesis and egg production in the fluke, which suggests a mutualistic relationship ^64^. Further studies are clearly needed to characterize how other viruses affect trematode phenotypes.

For fluke viruses to contribute to diseases and malignancy during human infection with the flukes, the virus(es) would have to be shed by the fluke and subsequently encounter human cells. We provide evidence that ORV and OSV are actively replicating in tegument cells at the surface of the adult parasite and that viral RNA is excreted with EVs. It is possible that the EV purification process resulted in co-enrichment of virions, because EVs and viruses tend to share density and size features that make them difficult to distinguish. Virions may have been co­enriched during EV purification due to the similar size and density profiles shared by EVs and viruses. Moreover, the expected size-range of OSV (25-120 nm diameter of Martellivirues) and ORV (a bullet shape with a length of up to 430 nm) agrees with a co-purification of virions together with sEV and lEV, respectively. Further studies will be needed to ascertain that ORV and OSV genomes detected in EV fractions are enclosed in EVs or whether free virions co­purified with EVs. However, the presence of subgenomic mRNA of OSV and mRNAs of ORV in lEVs, and of vsiRNA of ORV and OSV in sEVs indicates that the fluke actively excretes viral RNA into vesicles that are then internalized by cholangiocytes. This “Trojan EV” strategy, whereby infected cells export viral-derived RNAs into naïve cells is increasingly recognized as a strategy for viruses to sustain infection and replication while avoiding the host immune system, which in turn supports maintenance of chronic infection ^65,66^. We attempted to assess virus transmission to parasitized hamsters but detected very low RNA quantities in tissue samples. Future studies focusing on the biliary tree or using imaging approaches to test for the presence of viral RNA or proteins in host cells might profitably address this aspect.

Next, we turned to serological approaches and observed that *O. viverrini* infection was associated with the induction of a strong humoral response toward fluke-borne viruses. Our results show that most parasitized persons are exposed to fluke-borne viral proteins. The prevalence of the fluke viruses ranged from 13% and 85 % in our analyses, with viral loads sometimes reaching 1E+05, 1E+06, and 1+08 OPLV, ORV and OSV RNA molecules per individual adult fluke, respectively. We might expect different prevalence estimates with a larger sample size, as well as variation in virus prevalence among populations of *O. viverrini*, as previously known for *S. solidus* ^32^. Yet, with a mean of about 75 worms per infected individual (worm burden in *O. viverrini* infected individuals ranges from 1-3000 worms) ^67^, we infer that most *O. viverrini* infected people host flukes that harbor viruses. This high rate of exposure to viral proteins explains why fluke virus serology efficiently detected individuals with *O. viverrini* infections. Together, these results suggest the presence of *O. viverrini* viruses in all patients diagnosed with liver fluke infection-associated hepatobiliary diseases.

Validation of the first of Koch’s postulates prompted us to investigate further the association between infection with these novel fluke-borne viruses and disease development. Periductal fibrosis of the biliary tract is a prominent histological feature of chronic opisthorchiasis and is associated with proliferation of epithelial cells ^68^. Fibrosis is significantly associated with CCA, and is targeted in screenings for CCA using abdominal ultrasonography ^69^. Previous studies showed that levels of antibodies to *O. viverrini* excretory/secretory products, such as *Ov-*TSP2, were not associated with APF ^56^. Our serological analyses confirmed this earlier finding. In contrast, plasma levels of the pro-inflammatory cytokine IL-6 are associated with an increased risk of APF and CCA in a dose-dependent manner ^56,70^. IL-6 is well known for its role in chronic inflammation and fibrotic lesions and is considered a contributor to the development of hepatobiliary diseases during and following *O. viverrini* infection ^11^. Using the same cohort in which APF-subjects were matched to APF+ subjects by age, sex, infection and nearest-neighbor status ^70^, we showed that IgG titers against all three *O. viverrini* fluke viruses were significantly elevated in these study participants with APF. Our analysis also revealed that IgG titers were lower in patients with CCA than with APF. Measures of IgG avidity, however, revealed increasing IgG avidity among APF-, APF+ and CCA cases. Because the maturation of IgG avidity depends on the dose of antigen, the chronicity of infection and the number of reinfections, the association of elevated avidity against viral antigens in APF+ and, more strikingly, in CCA cases suggests that diseased patients were either exposed to higher doses of fluke-borne viruses, suffered from chronic infection, or experienced repeated reinfections. Alternatively, pro-inflammatory factors, including IL-6, can drive affinity maturation processes to produce high-avidity IgG clones. However, IL-6 upregulation can also be triggered by viral infection, promoting virus survival, and exacerbating clinical disease. Critically, our current data cannot distinguish the direction of this interaction: it remains unclear whether the IL-6 surge is a consequence of viral or fluke infection, and whether upregulation of IL-6 promotes virus survival, drives antibody maturation, or exacerbates clinical disease. Further studies are necessary to determine the respective roles and interactions between IL-6 and fluke-borne viruses in disease development.

Our results suggest, further, that fluke viruses can be targeted for diagnostic applications for *O. viverrini* infections and early CCA detection. Serological screening have been developed to diagnose past and ongoing infections, with indirect ELISA using crude antigens or ES products of the liver fluke as the method of choice ^71–73^. Herein, we developed LuLISA tests and showed that serological investigations of *O. viverrini* infections can be improved by combining serological tests against parasite antigens and species-specific associated viruses. Similar to cholangiocarcinoma in the Western world, early-stage CCA is mostly asymptomatic, and most patients presenting at a hospital have late-stage disease and receive supportive care with very poor prognosis ^74,75^. The detection of individuals with an early-stage tumor, when curative surgery is feasible, requires ultrasonographic screening at a medical facility or using a mobile unit ^76,77^. There is a pressing public health need to facilitate and improve access to APF and early CCA screening to improve patient survival^78^. Our IgG avidity results showing a dose-dependent avidity of antibodies against fluke-borne viruses in patients with APF and CCA prompted us to assess the feasibility of using serological approaches to predict the risk of diseases. The random forest model distinguishes among APF-, APF+ and CCA phenotypes with high sensitivity and specificity. Together, these results suggest that IgG titers and IgG avidity against fluke antigens could be used to predict the risk of diseases in *O. viverrini* infected individuals as an alternative to imaging in remote areas.

Nevertheless, a limitation of our study is that, whereas all participants were confirmed to have a current, patent *O. viverrini* infection at enrollment and were excluded for co-infection with hepatitis B or C virus or for alcoholic liver disease, a complete lifetime reinfection history of each participant could not be reconstructed retrospectively, since reinfection is common in this hyperendemic region despite periodic praziquantel treatment. This uncertainty is mitigated for the APF+ subgroup, for whom serial post-treatment fecal examinations over a two-year follow­up period confirmed the absence of reinfection during that interval. We further note that IgG avidity, which increased with hepatobiliary disease severity in our cohort (Fig. 9b), is itself a function of antigen dose, infection chronicity and reinfection frequency^57^, and may therefore serve as an indirect correlate of cumulative or repeated exposure to fluke-borne viral antigens across the disease spectrum studied here. Further longitudinal studies should be conducted to assess the evolution of IgG titer and avidity over the course of disease development and confirm the predictive value of this assay. Also, comparable studies in *C. sinensis* and *S. haematobium*-endemic settings are needed to determine whether fluke-borne viruses represent a shared risk factor for disease across all three carcinogenic trematodes. We expect that this unexpected discovery will eventually inform early detection of fluke-associated cancer, facilitate therapeutic development, and improve patient prognosis.

The three trematodes *O. viverrini*, *C. sinensis* and *S. haematobium* all are classified as Group 1 carcinogens because there is sufficient evidence that infection with these helminth parasites is definitely carcinogenic in humans ^2^. Until now, excretory/secretory proteins of liver flukes have been found to modulate angiogenesis, drive cell proliferation, and initiate pro-inflammatory responses, processes that exacerbate cancer by compromising host barriers ^27,79^. Our findings suggest that fluke-borne viruses are covert stowaway passengers of liver fluke infection and that helminth-infected individuals are chronically exposed to fluke viruses that are shed by the long-lived parasite. In ongoing efforts to identify the causal factors of fluke-associated cancer, the discovery that fluke-borne viruses are associated with increased risk of disease represents a major conceptual departure from the status quo, shifting focus onto viruses of flukes. Viruses are responsible for 10-15% of the worldwide cancer burden and up to 20% of the cancer burden in neglected populations ^2^. Oncogenic viruses establish long-term persistent infection, and it is the cell-cycle manipulations used by these viruses to promote replication and transmission that disrupt the cellular barriers to cancer and eventually promote carcinogenesis ^27,80^. Our results suggest that parasitized individuals may be chronically exposed to fluke viruses for the 3- to 30-years lifespan of adult flukes and that immune responses differ according to disease status. Yet, this study does not prove a causal role of the viruses in cancer development. Several non-mutually exclusive hypotheses should be considered: fluke viruses may (i) alter the production of virulence factors by the fluke or (ii) modulate the host immune response to promote a pro-oncogenic environment or (iii) directly impact the human cell cycle and initiate cell transformation. Elucidating the respective contributions of the parasite, its viruses, and associated microbes in carcinogenesis will require additional functional studies.

Last, we argue that it is the infection with the fluke and its biome, that should be considered carcinogenic, rather than the fluke alone. Recent findings showed that in addition to viruses, *O. viverrini* also carries *Helicobacter pylori*, the only bacetrium classified as a group 1 carcinogen ^81–84^. Indeed, there is a strong positive relationship between the intensity of liver fluke infection and the likelihood of hepatobiliary diseases and CCA, which may reflect increased exposure to fluke-borne viruses and other microbial symbionts at higher worm burden ^85^. Since the introduction of Koch’s postulates were introduced in 1884, the criteria for determining causality of a newly identified pathogen in disease development have evolved with our increasing understanding of microbial infection ^86–89^. We now face the challenge of determining the potential pathogenic roles of viruses (and bacteria) residing within flukes, parasites that are themselves Group 1 carcinogens.

## METHODS

### Animal Ethics statements

Male golden Syrian hamsters (*Mesocricetus auratus*) from the Animal Unit, Faculty of Medicine, Khon Kaen University, aged 6 to 8 weeks at the commencement of the study, are used throughout the study. Males were used to minimize estrous cycle-related variability in hepatic inflammatory/fibrotic responses and for consistency with priori characterization of this model. These hamsters were housed at five per cage, under conventional conditions, and fed a stock diet and water *ad libitum*. Their maintenance and care complied with the guidelines of the National Laboratory Animal Center, Thailand. Experimentation protocols were approved by the Animal Ethics Committee of Khon Kaen University according to the Ethics of Animal Experimentation of the National Research Council of Thailand, with the approval number IACUC-KKU-125/68.

### Human study participants

Participants were enrolled from a community-based cohort of opisthorchiasis-associated hepatobiliary morbidity in Khon Kaen Province, Northeast Thailand^56^. At enrollment, all participants underwent a structured clinical interview and physical examination; individuals with clinical, biochemical or ultrasonographic evidence of hepatitis B or hepatitis C virus infection-associated liver disease, or of alcoholic liver disease, were excluded from the study. Prior anthelminthic (praziquantel) treatment history was recorded for all participants by structured questionnaire at enrollment. Sex information was determined on the basis of self-reporting. A minority of participants (28.3%) reported prior praziquantel treatment. Given that reinfection is common in this hyperendemic setting, all participants were required to have a current, patent *O. viverrini* infection, confirmed by a positive fecal egg count (by formalin/ethyl acetate concentration technique), at the time of enrollment and serum collection, irrespective of prior treatment history. Following enrollment, all *O. viverrini*-positive participants received standard-of-care praziquantel (dose calculated according to body weight), with scheduled fecal examination and ultrasonography thereafter to monitor treatment outcome and reinfection status. APF-positive cases were treated with standard-of-care praziquantel (40 mg/kg) following ultrasonographic diagnosis and were followed for up to two years with serial fecal examinations; no *O. viverrini* eggs were detected during this follow-up period in any participant, indicating an absence of reinfection over this interval. All human study participants provided written informed consent using Informed Consents Forms approved by the Ethics Committee of Khon Kaen University School of Medicine, Khon Kaen, Thailand (reference number HE480528) and the Institutional Review Board of the George Washington University, Washington, D.C (GWU IRB# 020864).

### O. viverrini metacercariae, newly excysted juveniles, and adult worms

The metacercariae of *O. viverrini* were obtained from naturally infected cyprinoid fish from endemic area in Khon Kaen province, Northeast Thailand as described ^90^. Metacercariae were identified, collected under a dissecting microscope, and stored in saline solution at 4°C until used for hamster infections ^91^. Newly excysted juveniles (NEJs) of *O. viverrini* were obtained as detailed previously ^50^. Adult flukes were recovered from the livers of hamsters at ≥ 3 months after experimental infection with 50 metacercariae per hamster ^92^. In addition, urine, feces, serum, liver, gall bladder and kidney were recovered from five hamsters that had been infected with *O. viverrini* for one year and stored at -80°C until processing.

### RNA extraction, NGS library preparation and sequencing

Total RNA from six pools of 5 adult worms collected from two parasitized hamsters was extracted using the RNeasy plus mini kit (Qiagen, Valencia, CA, USA) following the manufacturer’s recommendation. RNA was eluted in 30 µl of water and 15µl of each sample was used to constitute the sequenced pool. RNA quality was evaluated using the RNA Pico chip (Agilent, Waldbronn, Germany) and an Agilent 2100 Bioanalyzer. The NGS library was constructed using the SMARTer Stranded Total RNA-seq kit v3-Pico input mammalian kit (Clontech, Takara Bio, San Jose, CA, USA) according to the manufacturer’s instructions and validated using the High Sensitivity DNA Chip (Agilent) and Bioanalyzer. DNA output of the library preparation was quantified with the dsDNA high-sensitivity kit (Invitrogen, Waltham, MA, USA) onto a Qubit 2.0 Fluorometer (Invitrogen). Sequencing was carried out on an Illumina NextSeq 2000 sequencer in a paired-end 2 × 100 bp format to achieve approximately 50 million paired reads per library.

### Data mining

*S. haematobium Illumina* RNA sequencing raw data were downloaded from NCBI SRA Bioprojects PRJEB32839 “The stage and sex specific transcriptome of the human parasite *Schistosoma mansoni*”, PRJEB37529 “Low input approaches to generate high-quality reference genomes for parasitic flatworms” and PRJNA78265 “*Schistosoma haematobium* Genome Sequencing and assembly”. *O. viverrini* Illumina RNA sequencing raw data were downloaded from NCBI SRA Bioprojects PRJEB65605 “ Transcriptome of *Opisthorchis viverrini* during intra-mammalian development”.

### Virus discovery

Raw sequencing reads were processed with Microseek, an in-house bioinformatics pipeline ^93^ that includes quality check and trimming, read normalization, *de novo* assembly, prediction of open reading frames (ORF), and taxonomic assignation of contigs and singletons using firstly an exhaustive and curated viral sequence database RVDB-prot ^94^, itself derived from the nucleic Reference Virus DataBase RVDB ^95^, and secondly a generalist NCBI/nr/nt databases.

### Viral genome completion

The draft viral genomes were reconstructed from the output of Microseek, with final manual check. Coding regions were annotated by homology to closely related viruses and domain identification. Consistency and absence of stop codon were checked. RNA extracts from the pool of adult worms used for virus discovery were employed for PCR to close genome gaps in the Opisthorhabdovirus. Briefly, an RNA extract from the pool of adult worms was retrotranscribed using random hexamers and the SuperScript IV Reverse Transcription system (Invitrogen) according to the manufacturer’s instructions. Primers flanking the gaps in the genome (Table S1) were designed, PCR was conducted with the Phusion high fidelity Taq polymerase (NEB, Ipswich, MA) according to the manufacturer’s instructions, and gel electrophoresis was performed to confirm the amplification. All PCR products were Sanger sequenced by Eurofins company.

We developed additional approaches aiming at identifying a putative third segments (segment M encoding for a Glycoprotein) of the Opisthorphenuilivirus, as expected based on close phylogenetic relatedness to *Phenuiviridae*. A first strategy was to identify panhandle sequences, hypothesizing that the segment M panhandle sequences of the Opisthorphenuilivirus would not be too distant from the sequences flanking its L segment (start ACACACAGACACCAG and end TGGTGGTCTTTGTGT) and S segment (start ACACACAGACACCCA and end GGGTGGTCTTTGTGT). Note that those flanking sequences differ by two nucleotides at the 5’termini and by only one at the 3’ termini. Therefore, we searched for both exact matches and near matches with one or two nucleotide variations. To validate that panhandle sequences could serve as filters for retrieving proteins from our data, we conducted a BLAST search on sequences containing the degenerate panhandle sequences against known S and L Opisthorphenuilivirus sequences, successfully retrieving fragments of both genome segments. The other selected sequences were then translated, and common sequences were aggregated and counted, before manual analysis to assess whether they encode putative glycoproteins.

The second approach did not rely on panhandle identification but, rather, it focused on similarities with known segment M sequences from other Bunyavirus species. A curated list of 18 representative segment M sequences was aligned using MAFFT. Conserved segments were extracted, and the most conserved loci were converted into regex expressions used to filter the sampled data. Out of nearly 100 million contigs and singleton segments, 180,000 passed the filtering, consisting primarily of short singletons. These were further filtered by selecting only those with bit scores higher than 30 when blasted against the curated list. Both methods employed relaxed constraints to allow the detection of sequences distant from known panhandle and segment M sequences. Unfortunately, this resulted in the selected sequences being predominantly short singletons or too distantly related to confirm their nature conclusively.

The third approach consisted in constructing a local BLAST database comprising all contigs and singletons generated from the de novo assembly of our NGS datasets. Reference M segment sequences from members of the Bunyavirus family were retrieved from public databases and translated into their corresponding protein sequences. These protein sequences were then used as queries in relaxed tBLASTn searches against our custom nucleotide database, allowing detection of highly divergent homologs at the amino acid level. This tactic increases sensitivity and enables identification of distant evolutionary relationships. No significant hits were detected, even under relaxed similarity thresholds confirming the absence of a M segment.

### AmpliSeq complete genome sequencing

Primer panels for multiple PCR were selected using PrimalScheme (PrimalScheme.com) using the genomes of ORV, OPLV and OSV as reference and an estimated amplicon length of 400bp (Table S2-S4). cDNA was obtained using the RT SuperScript IV first strand cDNA synthesis from 1 ng of *O. viverrini* RNA extract used for virus discovery. The PCR was conducted separately using pool 1 and pool 2 primers sets using the Q5 High Fidelity DNA Polymerase with a 30 sec denaturation step at 98°C, 35 amplification cycles (10sec at 95°C, 20sec at 58°C and 20sec at 72°C) and a final extension step of 3 min at 72°C. After clean up using AmpPure XP beads, the amplicons were quantified using Qubit kit DNA high sensitivity, and electrophoresis through agarose gel. Libraries were prepared using the Illumina TruSeq DNA PCR-free low throughput library prep kit following the manufacturer’s instruction and sequenced on a NextSeq 2000 with 600 cycles (2x300bp). Reads were aligned to the draft genomes, and the viral genomes were manually completed.

### Viral genome characterization and phylogenetic analyses

Open reading frames were predicted using NCBI ORF finder. Annotation of domains was deduced from comparisons against the Conserved Domain Database as implemented by BLASTp against the nr protein database to draw the genome composition (Fig. 1). Detailed information on the viral genomes is provided in Table S5. Initial supergroup assignments were determined from best BLAST matches. Predicted RNA-dependent RNA polymerase RdRp sequences from fluke-associated viruses, and representative sequences from related viral families and genera ratified by the ICTV and from recent metatranscriptomic studies, were aligned using the E-INS-I algorithm implemented in the program MAFFT version 7 ^96^. Next, all ambiguously aligned regions were removed using TrimAl version 1.2 implemented in Galaxy version 1.8.1^97^. Phylogenetic trees were inferred using the maximum likelihood method implemented in PhyML-SMS version 3.0 that includes Smart-Model selection ^98^ using the best-fit model and best of NNI and Subtree Pruning and Regrafting branch swapping. Support for nodes on the trees was assessed using an approximate likelihood ratio test with the Shimodaira–Hasegawa-like procedure. Phylogenetic trees were annotated using iTol (version 7.2).

### Total RNA and Small RNA Sequencing of EV

Total RNA and small RNA sequencing data described in ^40^ were re-analyzed for identification of viral sequences. Total RNA sequencing data were processed using Microseek ^93^. Clean reads were mapped against the reference viral genomes using CLC Genomics Workbench (Qiagen). Consistency and absence of stop codon were confirmed. The adapter was trimmed from the small RNA sequences using Cutadapt (version 4.9) and those reads where the adaptor was not found were discarded (parameters ‘-a AACTGTAGGCACCATCAAT --discard-untrimmed’). Trimmed reads in the 17-29 bp size range were mapped against ORV, OSV and OPLV using Bowtie2 (version 2.5.4) ^99^with parameters ‘-L 6 -i S,1,0.8 -N 0 --end-to-end’. Further processing was performed under Python (version 3.12.9) using the ‘bam2fastq’ and ‘make_read_stats’ functions provided by the libreads library (available at <https://gitlab.pasteur.fr/bli/libreads>). Reads mapping on either the forward or reverse strand were extracted from the alignment bam file using the pysam (version 0.23, wrapping samtools version 1.12 ^100^) library. Then, read size and composition statistics were computed, including composition at read start for each read size. The above steps (read pre-processing, mapping, extraction and statistics computation) were implemented as a Snakemake ^101^ (version 9.1.1) workflow available at https://gitlab.pasteur.fr/bli/19578_dheilly.

### PCR assays to assess virus presence

Samples from discrete developmental stages of *O. viverrini* were used to assess the presence of the virus through the parasite life cycle. Cercariae were shed from *Bithynia siamensis goniomphalos* snails obtained from endemic areas in two biological replicates. NEJs, representing the infective stage, were collected in six biological replicates, with 250 NEJs per replicate. Sixty-two individual adult worms from four hamsters were assayed. RNA extraction was performed for all parasite stages using Trizol reagent according to the manufacturer’s instructions (Invitrogen, USA). These RNAs were treated with DNAse I, using the RNase-Free DNase Set following the supplier’s instructions (QIAGEN). To ensure the highest sample quality, following DNase treatment, RNA were further purified using the RNeasy MinElute Cleanup kit, in accordance with the manufacturer’s instructions.

Total RNAs were reverse transcribed to cDNA using the RevertAid First Strand cDNA Synthesis Kit (Thermo Scientific, USA). Quantitative real-time PCR (qRT-PCR) was carried out using LightCycler 480 (Roche Diagnostics, Germany) and SYBR Green I (Roche) as the fluorophore. Primers were designed to detect either *C. sinensis* or *O. viverrini*-associated viruses through RT-qPCR. Primer sequences are provided in Table S6. Each RT-qPCR reaction, performed in triplicate, contained 10 µl of Maxima SYBR Taq (2×), 0.6 µl of each 10 mM primer, 2 µl of first-strand cDNA, and sterile water to a final volume of 20 µl. The PCR cycling conditions were initial denaturation at 95°C, followed by 40 cycles of 95°C for 5 seconds, 60°C for 30 seconds, and 72°C for 1 second. The *O. viverrini* actin gene (GenBank *EL620339*.1) was used as the internal control.

To assess virus presence in fluke infected hamsters, we used the RT-qPCR systems described above. Briefly, total RNA from hamster tissues were extracted using the Maxwell RCS Tissue kit (Promega) and retrotranscribed with random hexamers as described previously. SYBR Green qPCR assay FastStart Essential DNA Green Mastermix, Roche was performed in duplicates. PCR products were sized by electrophoresis through agarose gels, stained with SYBR green, purified with Qiagen PCR Clean-up kit, and sequence and specificity confirmed by Sanger sequencing.

### Viral RNA quantification

To quantify viral RNA within samples, we generated synthetic RNAs for the three viruses based on the sequences of ORV, OPLV and OSV. Briefly, primers flanking the region of the qPCR were designed, and the sequence of the T7 promotor was added in 5’ of the forward primer to allow *in vitro* transcription of PCR products (Table S7). Amplifications of portions of viral genomes was obtained from the pool of adult worms, purified with the PCR & Gel cleanup kit Macherey-Nagel, and *in vitro* transcribed using the Ambion^TM^ MEGAshortscript T7 Transcription Kit Invitrogen, according to the manufacturer’s recommendations. After purification, the number of genome copies was assessed by quantifying the concentration of RNA with the following formula : N= [C μg/mL ARN.10^-3^ x 6.023.10^23^ molecules/mol] / [length x 340g/mol.10^6^]. Synthetic RNAs were retrotranscribed as described above and 10-fold serial dilutions were used to determine the Limit of detection of the system. We quantified viral RNA within a total of 16 individual adult flukes positive to at least one virus. We conducted RT-qPCR as described above with a standard curve of the synthetic RNA. Viral RNA quantity was determined by converting the mean Ct value per samples into the mean viral RNA copy / PCR reaction.

### RNAscope in-situ hybridization to localize viral RNA

RNAscope® ISH (*in situ* hybridization) was used to localize the ORV, OSV and OPLV viral genomes and transcripts in adult *O. viverrini* worms. Adult worms from hamsters were washed thoroughly in 0.9% NaCl solution before fixation in 10% neutral buffered formalin (NBF) for 48h at room temperature, after which they were embedded in paraffin . For each virus, a panel of sense and antisense probes were developed and manufactured by Bio-Techne (https://www.bio-techne.com/; Minneapolis, MN, USA) (Table S8). Three µm sections of formalin-fixed, paraffin-embedded (FFPE) liver tissue were mounted on positively charged SuperFrost plus slides (Fischer Scientific). The assay was performed using an RNAscope ™ Multiplex Fluorescent Reagent Kit v2 (323100-USM, ACD®; Bio-Techne) using the recommended protocol. The samples were subjected to a protease-based pretreatment. The slides were then incubated with different RNAscope® 2-plex probes: a negative control probe (#320871), a positive control probe (Mau-Ppib-C1, #890851-C1), and target *O. viverrini* virus probes. After three amplification steps, fluorophores were added: TSA Vivid 570 (# 323272) to probe C1 and TSA Vivid 650 (# 323273) to probe C2, after which they were counterstained with DAPI. Slides were mounted, and imaged and analyzed using a digital slide scanner (AxioScan 7 digital slide scanner and Zen software (version 3.9.101) (Zeiss®, Oberkochen, Germany).

### Preparation of recombinant viral and fluke antigens

The ORV nucleoprotein, OSV capsid-core protein and OPLV nucleoprotein were selected as viral target antigens while the *O. viverrini Ov-*TSP-2 was selected as the parasite target antigen. Antigens have been synthesized with a carboxy-terminal His6-tag with Nco I and Xho I restriction site at each end, respectively (GeneArt^TM^, Thermo Fisher). The synthetic genes were subcloned by cleavage of pET-23 plasmid and expression gene by Nco I and Xho I (NEB), ligated into the plasmid (T4 ligase, NEB). Gene sequences were validated by Eurofins using T7 and T7 terminator. T7 Express LysY/Iq *E. coli* strains were chemically transformed with the pET-23/viral antigen constructs and plated in Petri dishes on LB agar supplemented with carbenicillin 100 µg/mL. Erlenmeyer flasks containing 300 mL of NZY medium (NZ-Biotech) and carbenicillin 100 µg/mL were seeded from single colonies and bacteria were grown, orbitally agitated at 180 RPM overnight at room temperature (RT). The liquid cultures were centrifugated 10 min at 4000 RPM (4-16KS, Sigma) in 50 mL conical tubes (ThermoFisher). Supernatants were discarded and pellets were resuspended in 2x PBS, 20mM phosphate, 20 mM imidazole, 300 mM NaCl, lysozyme 30 mg/mL, Triton 0.1%, EDTA-free protease inhibitor 1x (Roche). B-PER™ II Bacterial Protein Extraction Reagent 1X (ThermoFisher Scientific). The suspensions were homogenized and left for 15 minutes at RT to enable lysis. The lysates were placed on ice before addition of 50 µL of RNAse and DNase I at 10 mg/mL (Roche), and MgCl_2_ (Sigma-Aldrich) at 2 mM final concentration. The lysates were clarified by centrifugation, 30 min 4 °C, 13300 g (5427R, Eppendorf). For each antigen, a 5 mL His-Trap column (Cytiva) was preequilibrated with 25 mL 2x PBS, 20 mM imidazole using an AKTA pure liquid chromatographic system (Cytiva). The supernatant was loaded on the column 4 mL/min, then washed with 65 mL 2x PBS, 20 mM imidazole. Proteins were eluted with 2x PBS, 500 mM imidazole. The 280 nm-absorption was recorded in real time, and samples were collected in 1 mL fractions. High absorption fractions were separated by electrophoresis through stain-free 4­15% acrylamide-bisacrylamide SDS-PAGE gels (Bio-Rad, Hercules, CA) in SDS 10% (Sigma-Aldrich). Fractions containing recombinant proteins were pooled and concentrated to a volume of 0.5 mL by centrifugation using 30 kDa-cutoff filtration devices (Ultra 30k, Amicon). A Superdex 200 column (Cytiva) was preequilibrated using 50mL PBS at 0.8 mL/min (Akta pure, Cytiva). The 0.5 mL fraction was loaded on the Superdex 200 column at 0.8 mL/min for a size­exclusion separation and 0.5 mL fractions were collected. High absorption fractions were electrophoresed on stain-free 4-15% acrylamide-bisacrylamide gels (Bio-Rad) in SDS 10% (Sigma-Aldrich). Fractions containing proteins were pooled and EDTA-free protease inhibitor 1x (Roche) was added for storage at 4°C and -80°C. Protein concentrations were measured at 280 nm.

### Serologic Luciferase-Linked Immuno-Sorbent Assay LuLISA for detection of IgG

White 96-well plates (FluoroNunc C96 MaxiSorp™, Thermofisher) were coated with the recombinant antigen at 1 µg/mL in carbonate buffer pH 9.3, 50 µL/well, overnight at 4°C. The plate was washed 3 times with 100 µL of PBS Tween 0.1% per well using the portable aspiration system Vacusip (Integra). Sera were diluted 5 times in glycine-HCl buffer pH 1.8 for acid immun-complex dissociation (ICD) at RT for 90 mins. ICD were neutralized with Tris-HCl buffer pH 9.0 to reach a final pH of 7.5. Immediate dilution was performed in PBS 0.1% Tween20 buffer to reach a final serum dilution of 1:150. Diluted sera were loaded at 50 µL/well, incubated for 60 min at RT then washed 3 times with 100 µL of PBS, 0.1% Tween 20 per well. An anti-hamster IgG Fc-specific VHH (variable heavy chain domain, or nanobody) or an anti­human IgG Fc-specific VHH, expressed as a chimera with the luciferase JAZ (described in the patent applications Rose et al., Luciferase linked immunosorbent assay, EP4143306A1, US20230145894A1, CA3176386A1), was added to wells: 50 µL / well, 1 ng/mL of VHH-JAZ in PBS Tween 0.1% incubated 1 hour at room temperature. The plate was washed 3 times with 100 µL of PBS Tween 0.1% per well. The luciferase substrate, Hikarazine Q108 ^102^ was loaded at 50 µL / well, Q108 13.5 mM in PBS. The bioluminescence was measured extemporaneously using a luminometer (Centro, Berthold Technologies, Bad Wildbad, Germany) at 0.5 s / well ^52^. Avidity Index determination ^57^ was performed on 72 samples of each group of subjects (APF-, APF+, CCA). Each sample was assayed by LuLISA with two replicates on the same plate. After the serum incubation test at dilution 1:300 (ICD was performed), the plates were washed once with PBST 0.1% and then one replicate was incubated with PBST 0.1% and the other with PBST 0.1% + 6M urea for 10 minutes at RT. After this step, the plates were washed two times in PBST 0.1%, and the following steps were the same as for the LuLISA described earlier. The Avidity Index (AI) was determined as follow: AI (%) = RLU (urea)/ RLU (PBST) *100.

### Statistical analyses

The antibody titer was quantified as RLU/s (Relative Light Unit per second) and expressed as RLU/s or RU (Relative Unit). The RU was determined through a LuLISA standard curve in which a serial dilution of a known concentration of specific IgG was performed. Each standard curve was approximated with a 4-parameters Hill model using the package minpack.lm of RStudio (R version 4.3.0). To normalize the data among plates, two positive samples were assayed on each plate and used for calibration.

The usefulness of serological tests for diagnosis of opisthorchiasis in terms of sensitivity, specificity, and positive predicted values and negative predicted values was estimated by Receiver Operating Curve (ROC) analysis using R package pROC and Youden criterion to determine optimal cutoff. Bootstrapping was performed to evaluate metrics and their 95% confidence intervals. Statistical analyses and graphics generation were performed on RStudio (R version 4.3.0). Kruskal-Wallis test for difference in distributions was conducted for each antibody response (*Ov-*TSP-2, ORV nucleoprotein, OPLV nucleoprotein, OSV capsid-core protein). Reported effect size was calculated as ɳ^2^_H_ = (H - k + 1)/(n - k); p-values were adjusted using Benjamini-Hochberg correction. Post hoc pairwise comparisons were performed using two-sided Wilcoxon-Mann-Whitney tests. Reported effect size was calculated as r=Z/√n, where n is the total number of observations; p-values were adjusted using Benjamini-Hochberg (BH) correction to account for multiple comparisons ^103^.

The percent distribution of selected demographic characteristics was calculated for (1) parasitized by *O. viverrini* and APF negative (n=184) and (2) parasitized by *O. viverrini* and APF positive (n=184). The estimated sample size for the unmatched case-control study was 85, providing 80% power at a 95% confidence level 2-sided with a significance level of *p* < 0.05 with a Bonferroni correction for multiple testing. Cases were matched by age (10 year age intervals), sex and nearest-neighbor status (same village) ^70^. Adjusted odds ratios and 95% confidence intervals for log transformed antibody titers RU and their association with APF were determined using age, sex and liver fluke eggs per gram of feces (EPG) adjusted multiple logistic regression analyses. Confidence intervals for quartiles of RU and their association with APF were determined using age and sex adjusted logistic regression analyses.

We used the random forest method to train classification models predicting either infection status (French non endemic subjects vs Thai infected subjects) or diagnostic (FR vs APF-vs APF+ and CCA vs APF-vs APF+). Classification predictions were made based on antibody titers from all viruses, along with their avidity measure for models including CCA patients. To best estimate models performances, we defined a training set (n=385) and an independent validation (n=170, extended data 2) set, using covariate-adaptative randomization to make sure both were homogeneous and representative of the whole cohort regarding viral titers, avidity, infection status and diagnostics. Training set was used for the training and tuning phase of all models, using grid search and 10-fold cross validation to select best performing parameters (node size and number of tree) while limiting over-fitting. AUC (ROC metric in caret ^104^) was used for tuning as it is less sensitive to class imbalance as accuracy. Validation set was then used for performance evaluation of the final models. Training and evaluation were both performed on R 4.4.2 using caret package ^104^.

## DATA AVAILABILITY

The genomes of viruses of *O. viverrini* and *S. haematobium* has been assigned Genbank accession numbers PX443621 to PX443624 and BK075116 to BK075119, respectively. Sequencing data of *O. viverrini* used for virus discovery and viral genome completion have been submitted under BioProject ID PRJNA1337339. Sequencing data of *O. viverrini* extracellular vesicles miRNA and mRNA have been submitted under BioProject ID PRJNA1353125. All the source data, phylogenetic trees, sanger sequences and microscopy images are available on Zenodo (10.5281/zenodo.22128439).

## CODE AVAILABILITY

hyphy 2.5.31; PhyML & MAFFT were implemented through NGPphylogeny https://ngphylogeny.fr/; MICROSEEK is an in-house pipeline that uses ALIENTRIMMER v. 2.0 https://gitlab.pasteur.fr/GIPhy/AlienTrimmer for read trimming / clipping, BBNORM from BBMAP v.39.06 package https://sourceforge.net/projects/bbmap/ for coverage normalization, MEGAHIT v.1.2.9 https://github.com/voutcn/megahit for assembly, an in-house ORF-finder https://figshare.com/articles/code/translateReads_py/7588592, and then DIAMOND v. 2.1.9 https://github.com/bbuchfink/diamond/ and NCBI BLAST v. 2.16.0+ ftp://ftp.ncbi.nlm.nih.gov/blast/executables/blast+/2.16.0/ both for sequence searching. All newly generated code related to the small RNAseq analysis and statistical analyses of serology data is available on Zenodo (10.5281/zenodo.22128439).

## ACKNOWLEDGMENTS

We thank Giovanni Begliomini, Hugo Boutonnet, Philippe Pérot, Souand Mohammed Ali, Delphine Chrétien, and Youna Maillié for providing technical support. We thank Chloé Baum, Biomics Platform, C2RT, Institut Pasteur, Paris, France, supported by France Génomique (ANR-10-INBS-09) and IBISA. We thank Caroline Demeret for her thoughtful comments on an earlier version of the manuscript. This work was supported by a project Emergence Flash from the Department of Research Applications and Industrial Relations (DARRI) of Institut Pasteur, Paris, France to NMD, by the Comité de Paris de la Ligue Contre le cancer Grant RS25/75-30 to NMD, and by the Agence Nationale de la Recherche project ANR-2A-MRSEI-2023 to NMD, BS, PJB and AL. TRose reports support from the French public bank investment (BPI France), Ile-de-France grants (SESAME, project Bioassays), and Institut Carnot MS grants (ArThy Emergence and Maturation). In addition, SC, TL, AL and PJB received support from award RO1 CA164719 from the National Institutes of Health, National Cancer Institute (SA) and SC received grant from Faculty of Medicine Khon Kaen University (Grant Number IN69061). BS is a Senior Research Scholar supported by Khon Kaen University. The funders had no role in study design, data collection and analysis, decision to publish, or preparation of the manuscript.

## AUTHOR CONTRIBUTIONS

NMD conceptualized the study. NMD, BS, TRose, TL and DHardy supervised the experimental work. SC, TR, AF, DH, ST, SG and NMD conducted the experimental work. SC, TR, ST, AF, DH, EJ, TB, BL, JK, LC, JS, TRose, BS, and NMD participated in data curation and formal analysis of data. TR, EJ and BS conducted statistical analyzes. BS collected patient samples and clinical data. SC conducted animal experiments and managed samples access and processing. JS, TL, TRose, PJB, and AL provided key resources necessary to the project, participated in data interpretation and manuscript preparation. SC, TR, AF, DH, EJ, BL, JK, BS, NMD wrote the first draft of the manuscript. SC, TRose, AL, PJB, BS, NMD ensured funding acquisition. SC, TR, AF, DH EJ, BL, and NMD prepared figures. All authors reviewed and improved the manuscript.

