## supplementary figures and tables for "Fluke-borne viruses are a risk factor and diagnostic target for diseases caused by carcinogenic trematodes"

### Appendix

Sujittra Chaiyadet, PhD[1,2]†, Tom Roblin, MSc[3]†, Anais Fauchois, MSc[4,5]†, Sarah Temmam, PhD[4,5]†, David Hing, BSc[6], Elise Jacquemet, MSc[7], Thomas Bigot, PhD[4,5,7], Blaise Li, PhD[7], Julia Kende, PhD[7], Lisandru Capai, PhD[4,5], Javier Sotillo, PhD[8], Thewarach Laha, PhD[9], Sophie Goyard, PhD[3], David Hardy, PhD[6], Thierry Rose, PhD[3], Alex Loukas, PhD[10], Paul J Brindley, PhD[1,11], Banchob Sripa, PhD[1,2], Nolwenn M Dheilly, PhD[4,5]*

**Table of content**

Fig. S1.2
Fig. S2. 4
Fig. S3.4
Table S1.5
Table S2.6
Table S3.8
Table S4.10
Table S5.1
Table S61
Table S8.3
Table S9.4
Table S10.5
Table S11**Erreur ! Signet non défini.**
Table S12.8
Table S13.7
Table S14.9
Table S15.10
Table S16.11
Table S17.12

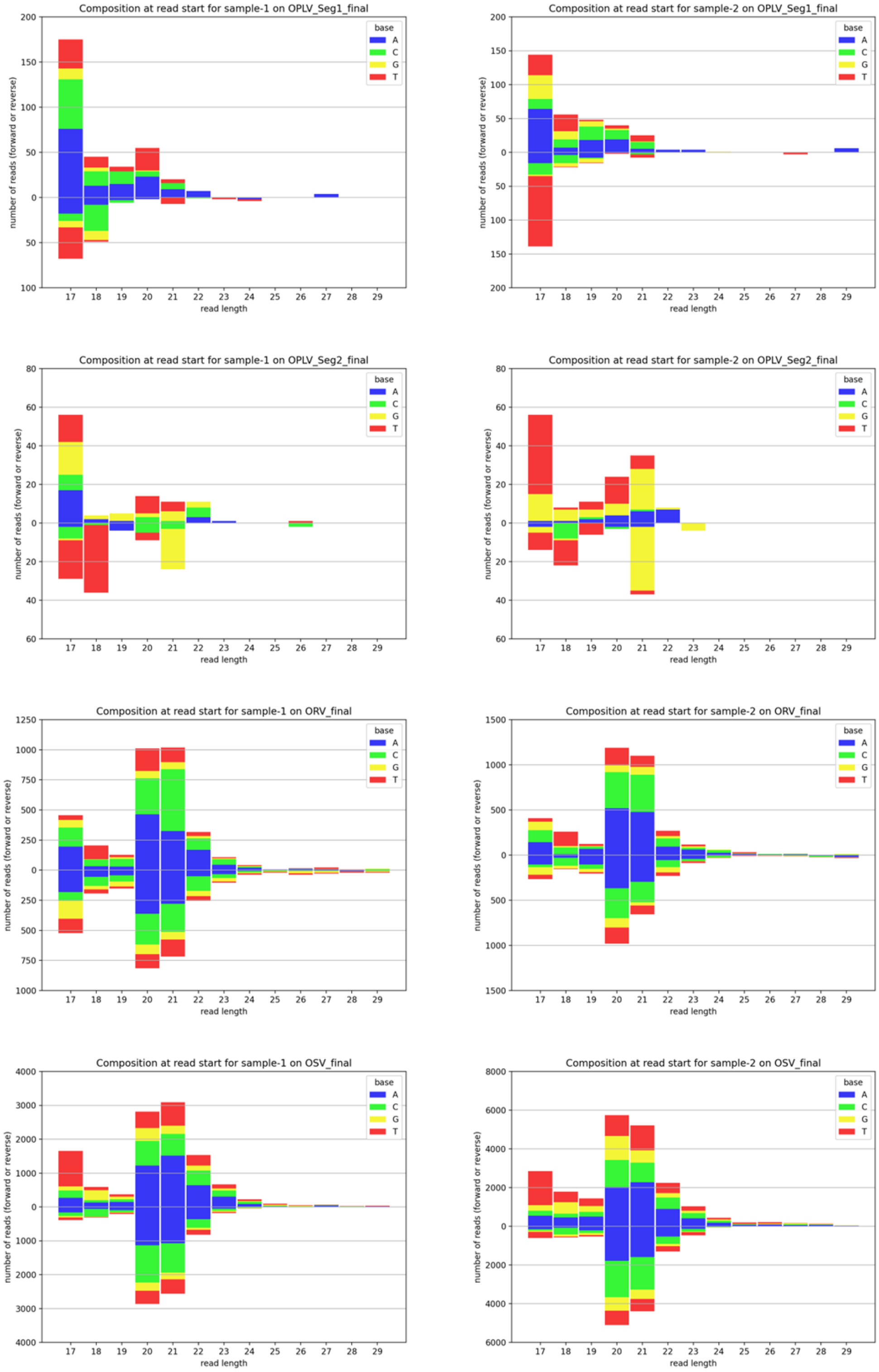

**Fig. S1.**

Size distribution, polarity and the 5-terminal nucleotide of total viral small RNA from two EV libraries.

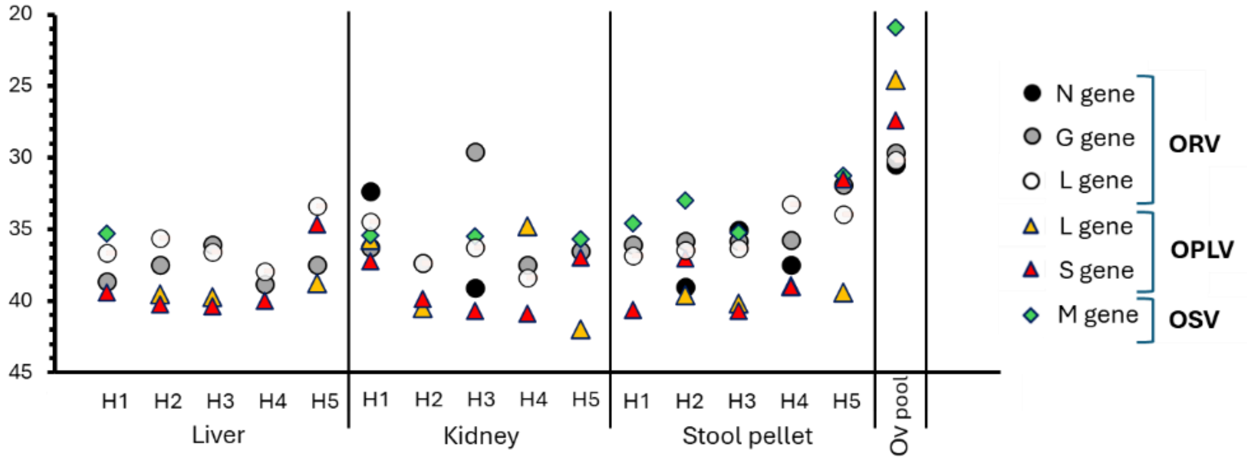

**Fig. S2. Molecular detection of fluke viruses in parasitized hamsters** detection of ORV, OSV and OPLV nucleic acid in the liver, kidney and stool pellet of parasitized hamsters. The pool of *O. viverrini* samples used for virus discovery was used as positive control

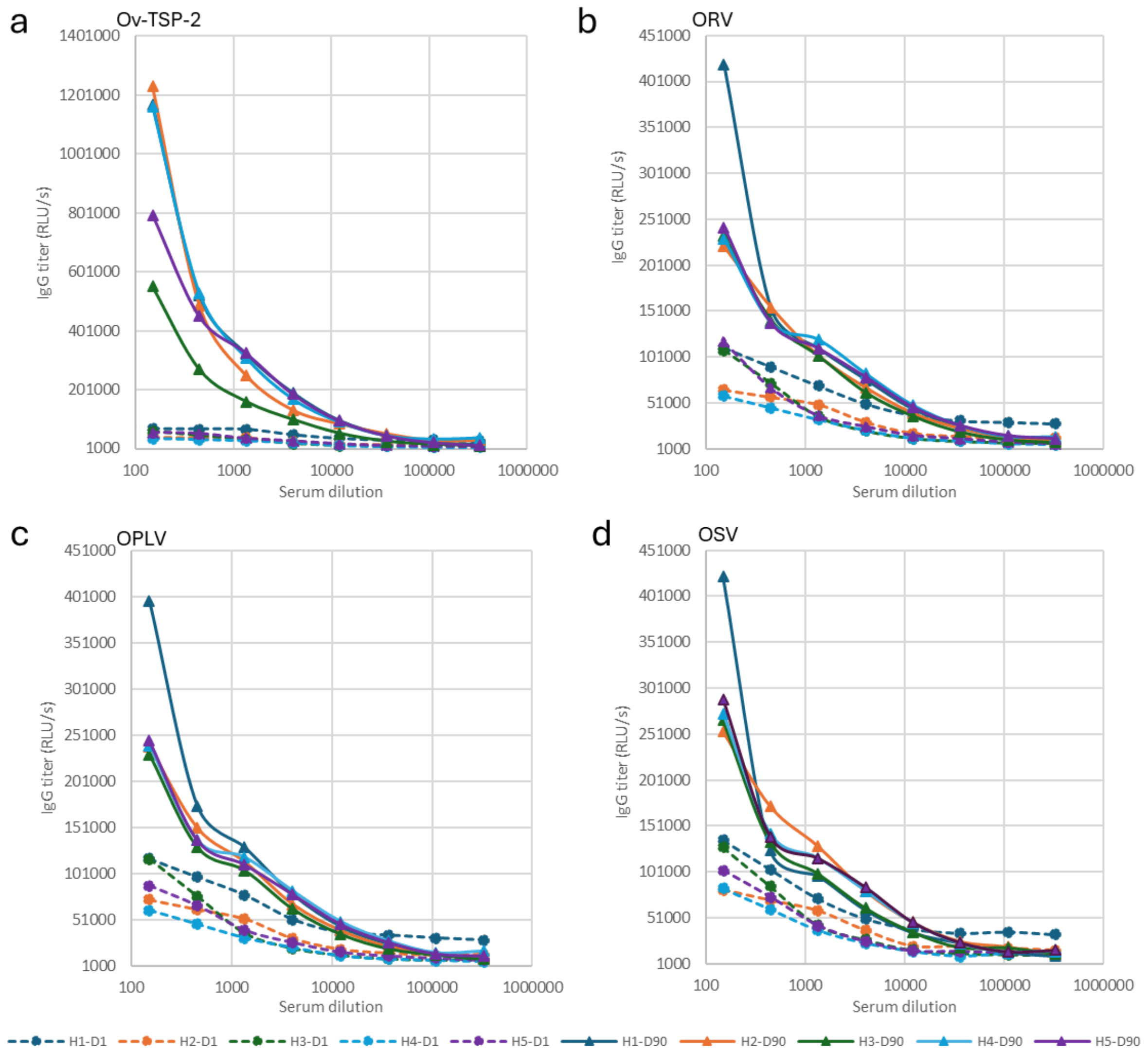

**Fig. S3. Serological detection of fluke viruses in parasitized hamsters** (c-f) The antibody titer in hamsters was determined as the last dilution in which the ratio of signal between the day 90 post-infection (D90) sample and the first day post-infection (D1) sample was superior to or equal to 2.0. The figures display the results obtained for Hamsters 1 to 5 (H1 to H5) for c) *Ov*-TSP-2, d) ORV, e) OPLV et f) OSV.

**Table S1.**
Primers used for Opisthorhabdovirus genome completion

| OligoName | Sequence | Comment |
|---|---|---|
| Oviv_rhabdo_F1 | CGATCGTGAACATACTCAGAATGT | PCR annealing 60°C - 413 pb |
| Oviv_rhabdo_R1 | GATCAGGCGTGATATTGAGCTGA | |
| Oviv_rhabdo_F2 | GACTCTCACMCGACCCTGG | PCR annealing 60°C - 699 pb |
| Oviv_rhabdo_R2 | GAAGATKGGGAAAGGGGTCTC | |
| Oviv_rhabdo_F3 | CTCAAACACCGATGTCCTCAC | PCR annealing 60°C - 504 pb |
| Oviv_rhabdo_R3 | CGATCTAAGACATGATTGATCTGG | |
| Oviv_rhabdo_F4 | ACCGTCAGCGGGTTGAACTC | PCR annealing 60°C - 521 pb |
| Oviv_rhabdo_R4 | CTCAGCRCAAAGTCCGAGAG | |
| Oviv_rhabdo_F5 | GATAATAATGAATGGGACCTTGTC | PCR annealing 58°C - 566 pb |
| Oviv_rhabdo_R5 | TGATCTAGGAAAGAGAAGACCC | |
| Oviv_rhabdo_F6 | AGTGGCATGGGAGCTTAACATGA | PCR annealing 60°C - 343 pb |
| Oviv_rhabdo_R6 | TGAYGCTGCTGATCCGAGGA | |
| Oviv_rhabdo_F7 | ACTGTCCCGTACCCGATTCA | PCR annealing 58°C - 192 pb |
| Oviv_rhabdo_R7 | ACCTTGAGAGTAGCGGAGCC | |
| Oviv_rhabdo_F8 | TGCACGATCCTGTTTGGATCG | PCR annealing 58°C - 540 pb |
| Oviv_rhabdo_R8 | ACTGTCGAYTCTTCTACCTTGG | |
| Oviv_rhabdo_F9 | GATTGTTGAGACCCCTCTATCC | PCR annealing 60°C - 498 pb |
| Oviv_rhabdo_R9 | AAGCTCGAACGCATTCGATCTG | |
| Oviv_rhabdo_SP1 | AGCTTTGAGGAGCTCAGCTGC | 5' RACE PCR annealing 58°C |
| Oviv_rhabdo_SP2 | TCCGGGAGTGTTCCCGTCA | |
| Oviv_rhabdo_SP3 | TGGRACATTCTGAGTATGTTCAC | |
| Oviv_rhabdo_SP5 | CAGTGCGTGAGCTATTCCGA | 3' RACE PCR annealing 58°C |
| Oviv_rhabdo_SP6 | GACCGCTGTGATATGCGACG | |

**Table S2.**

Primers used for complete genome sequencing of ORV.

| Name | Pool | Sequence | Size (bp) | %GC (%) | Tm (°C) |
|---|---|---|---|---|---|
| Rhabdo_1_L | 1 | CTCAACGATCGTGAACATACTCAGA | 25 | 44.00 | 61.05 |
| Rhabdo_1_R | 1 | GCTGATGGAGGTAGAAGATGACG | 23 | 52.17 | 60.80 |
| Rhabdo_2_L | 2 | GGGACCGGAATAGTTGAAGCAT | 22 | 50.00 | 60.60 |
| Rhabdo_2_R | 2 | ATAGGCGAATCATGATGACGGC | 22 | 50.00 | 61.11 |
| Rhabdo_3_L | 1 | GCTCCGACATCCTTAATGAGATCA | 24 | 45.83 | 60.58 |
| Rhabdo_3_R | 1 | TCATGAGATACATAAAGGGAACGGC | 25 | 44.00 | 61.12 |
| Rhabdo_4_L | 2 | TCGGCCTTACTTCCCCTACATT | 22 | 50.00 | 61.08 |
| Rhabdo_4_R | 2 | CTTCGCTCATGATCGTTGGTGT | 22 | 50.00 | 61.42 |
| Rhadbo_5_L | 1 | GTCATCAAACTGATCTGGAGGACC | 24 | 50.00 | 61.24 |
| Rhabdo_5_R | 1 | TTTTGTCCTGGGGAGGGAGTT | 21 | 52.38 | 61.09 |
| Rhabdo_6_L | 2 | GGACATTTCTGCCTCTGCGTAT | 22 | 50.00 | 60.92 |
| Rhadbo_6_R | 2 | TTTTGAGATTAGGCCAGCTGGG | 22 | 50.00 | 60.81 |
| Rhabdo_7_L | 1 | TCAACGATGCAACGAAGGAAGT | 22 | 45.45 | 60.99 |
| Rhabdo_7_R | 1 | TGATCGAGTCCGGCTTTAGTGA | 22 | 50.00 | 61.38 |
| Rhabdo_8_L | 2 | CCAAAATCGATCGTGCTTGAGC | 22 | 50.00 | 60.96 |
| Rhabdo_8_R | 2 | AGATGGGGAAAGGGGTCTCT | 20 | 55.00 | 59.59 |
| Rhabdo_9_L | 1 | TCCGTGATTGCTCTTTTAAAAACACT | 26 | 34.62 | 60.62 |
| Rhabdo_9R | 1 | ATGGTGTTACTGTAGGCCCCTT | 22 | 50.00 | 61.29 |
| Rhabdo_10_L | 2 | AGTGGAGCAAGCCTGGTGTA | 20 | 55.00 | 61.14 |
| Rhabdo_10_R | 2 | ATCAACTCGTGCATCCAGATGG | 22 | 50.00 | 60.93 |
| Rhabdo_11_L | 1 | CGTAGGCAACATACAAAAACCGC | 23 | 47.83 | 61.45 |
| Rhabdo_11_R | 1 | CAAAGAGACTCCTGGAAACTGCT | 23 | 47.83 | 60.75 |
| Rhabdo_12_L | 2 | TATTGTGGCCCTAGCGATTGC | 21 | 52.38 | 60.92 |
| Rhabdo_12_R | 2 | CGTCCTGTAGATAAGACATGATCCC | 25 | 48.00 | 60.72 |
| Rhabdo_13_L | 1 | ACTTAGGGCGCGAAAAGTGG | 20 | 55.00 | 60.97 |
| Rhabdo_13_R | 1 | TCGACTGAGGTAGATTCTACCCG | 23 | 52.17 | 61.00 |
| Rhabdo_14_L | 2 | TACCGTGTGGGCGATTGTTAAG | 22 | 50.00 | 61.11 |
| Rhabdo_14_R | 2 | AACTCATCCATCAACCGTTGGG | 22 | 50.00 | 61.06 |
| Rhabdo_15_L | 1 | GTCTGCGTGCGAAACTAGTCTT | 22 | 50.00 | 61.35 |
| Rhabdo_15_R | 1 | TCGTCGGTTTGTTCTCCTCTCT | 22 | 50.00 | 61.26 |
| Rhabdo_16_L | 2 | CGGGTTGAACTCAAAGAGCTGA | 22 | 50.00 | 60.99 |
| Rhabdo_16_R | 2 | TCAAGGTCTCCTGCTCGGAAT | 21 | 52.38 | 61.00 |
| Rhabdo_17_L | 1 | CATCGCTCAGCTGATACTACAGG | 23 | 52.17 | 60.86 |
| Rhabdo_17_R | 1 | AGCTCTGCCGTGTAAAATCCTG | 22 | 50.00 | 61.12 |
| Rhabdo_18_L | 2 | GCAACGAAGGATTTCAGCAGTG | 22 | 50.00 | 60.84 |
| Rhabdo_18_R | 2 | GCTGGACCTCTTGCTTGATGAA | 22 | 50.00 | 61.06 |
| Rhabdo_19_L | 1 | CTTACACGTCGGCACTACTCAT | 22 | 50.00 | 60.34 |
| Rhabdo_19_R | 1 | CTACTTGGTAAGCGCGCCTTAT | 22 | 50.00 | 60.98 |
| Rhabdo_20_L | 2 | CTGCCTTGGTTATTATCGCCGA | 22 | 50.00 | 60.99 |
| Rhabdo_20_R | 2 | TCTCTCTCCTTCTCACGTAGCC | 22 | 54.55 | 60.86 |
| Rhabdo_21_L | 1 | CCTCAGTTCAGGAAATCATGGCT | 23 | 47.83 | 60.88 |

| | | | | | |
|---|---|---|---|---|---|
| Rhabdo_21_R | 1 | CCCGATGAGAGTTAAGAAAAAGATGAG | 27 | 40.74 | 60.47 |
| Rhabdo_22_L | 2 | TCGACTTTGAGAAATGGAACCTCA | 24 | 41.67 | 60.52 |
| Rhabdo_22_R | 2 | CGAAACTTCATGAAGCTTGCGAC | 23 | 47.83 | 61.44 |
| Rhabdo_23_L | 1 | GGGACAACCAGGTAATAGCACTC | 23 | 52.17 | 60.94 |
| Rhabdo_23_R | 1 | AGGGATGATAAGCTCTTAAACAGCC | 25 | 44.00 | 61.07 |
| Rhabdo_24_L | 2 | CGACGAAACCTCTTCCATTCGA | 22 | 50.00 | 60.85 |
| Rhabdo_24_R | 2 | CCGGGATAAGATCGGGGAGTAA | 22 | 54.55 | 61.00 |
| Rhabdo_25_L | 1 | ACACAATCTTTGACCTGGACCC | 22 | 50.00 | 60.67 |
| Rhabdo_25_R | 1 | TGTTGTTCACGGTACATCAGGC | 22 | 50.00 | 61.31 |
| Rhabdo_26_L | 2 | CCCGCTTGATGTCTGACATTGT | 22 | 50.00 | 61.38 |
| Rhabdo_26_R | 2 | GAAACGGGCCGTGTTGAGATAA | 22 | 50.00 | 61.37 |
| Rhabdo_27_L | 1 | CCATAAGCCACCAACCTCACTC | 22 | 54.55 | 61.12 |
| Rhabdo_27_R | 1 | TGCAGTGTGTGGAGGATCGT | 20 | 55.00 | 61.49 |
| Rhabdo_28_L | 2 | GTGCGGAACATAGATTCGCAGA | 22 | 50.00 | 61.23 |
| Rhabdo_28_R | 2 | ATCGTCACGATTCAATCCTCCC | 22 | 50.00 | 60.40 |
| Rhabdo_29_L | 1 | ACCATCTTTCCCTCATATCCCCT | 23 | 47.83 | 60.64 |
| Rhabdo_29_R | 1 | TCAGTCCCAAGGATCGATACGA | 22 | 50.00 | 60.61 |
| Rhabdo_30_L | 2 | GTTTGGATTGGATCTTCGACATCG | 24 | 45.83 | 60.57 |
| Rhabdo_30_R | 2 | ACGGTTGGTGAGTGATAGAATGTC | 24 | 45.83 | 60.94 |
| Rhabdo_31_L | 1 | CTTTGTCTCGGCAGTTCGTGA | 21 | 52.38 | 60.97 |
| Rhabdo_31_R | 1 | ACCATGATGATGTCGGGTATCC | 22 | 50.00 | 59.88 |
| Rhabdo_32_L | 2 | ATGGTATCTCAGCAGCCCACT | 21 | 52.38 | 61.08 |
| Rhabdo_32_R | 2 | CTCTGCTGACTTGGTTAGATCACC | 24 | 50.00 | 61.24 |
| Rhabdo_33_L | 1 | GAGTTGCAACCCTCTGGACTTT | 22 | 50.00 | 60.93 |
| Rhabdo_33_R | 1 | CGAGCTGACACGTTTCGCTT | 20 | 55.00 | 61.56 |
| Rhabdo_34_L | 2 | CGATCAGTTTCTAGAAGCCGGTT | 23 | 47.83 | 60.93 |
| Rhabdo_34_R | 2 | GATTTTCAGACGATAAATAATGGACCCTC | 29 | 37.93 | 61.12 |
| Rhabdo_35_L | 1 | CACAACGATTGAAGCAAAGCTGA | 23 | 43.48 | 60.80 |
| Rhabdo_35_R | 1 | ATACTCAGCTCTTATCGTGCGG | 22 | 50.00 | 60.21 |

**Table S3.**

Primers used for complete genome sequencing of OSV.

| Name | Pool | Sequence | Size (bp) | %GC (%) | Tm (°C) |
|---|---|---|---|---|---|
| Opisthorsi_1_L | 1 | GGCTGGGAAACACGATCATACA | 22 | 50.00 | 60.86 |
| Opisthorsi_1_R | 1 | GGAAACGCTCTCACCAACTTCT | 22 | 50.00 | 60.99 |
| Opisthorsi_2_L | 2 | CTAAGCGTCGGGTTAGTAGTGATC | 24 | 50.00 | 60.87 |
| Opisthorsi_2_R | 2 | ATCTGACAACTTCCTGCATGCC | 22 | 50.00 | 61.39 |
| Opisthorsi_3_L | 1 | CGTTCAAGACGCTAATCGAAATGC | 24 | 45.83 | 61.60 |
| Opisthorsi_3_R | 1 | TCCGAAGCTCATGACAATAGTGC | 23 | 47.83 | 61.24 |
| Opisthorsi_4_L | 2 | CCGGAAGTATCATCGTGATGGG | 22 | 54.55 | 60.79 |
| Opisthorsi_4_R | 2 | GCACACTCGATTCTTTTAAGTCGG | 24 | 45.83 | 60.74 |
| Opisthorsi_5_L | 1 | GGTTGTGCCAAAGAAACTATGGG | 23 | 47.83 | 60.56 |
| Opisthorsi_5_R | 1 | TTCTTCGCCATCTTCACTGACC | 22 | 50.00 | 60.79 |
| Opisthorsi_6_L | 2 | GAAACAGACTTGGCGGAAGAGA | 22 | 50.00 | 60.73 |
| Opisthorsi_6_R | 2 | CCACTAGGATCGTGCGCAAT | 20 | 55.00 | 60.55 |
| Opisthorsi_7_L | 1 | GAATTGTGAAGATTCGCTGCGC | 22 | 50.00 | 61.52 |
| Opisthorsi_7_R | 1 | AACTTCCACCGAGAGTTGAGGA | 22 | 50.00 | 61.20 |
| Opisthorsi_8_L | 2 | GCACTGGTTGAGGAAATGATGC | 22 | 50.00 | 60.60 |
| Opisthorsi_8_R | 2 | AGAAGTTCTCGTACGAAAACAGAGT | 25 | 40.00 | 60.60 |
| Opisthorsi_9_L | 1 | GGGTTGTCGTCGGTCTTACAAA | 22 | 50.00 | 60.98 |
| Opisthorsi_9_R | 1 | AACCACAGACGAGAACTTCTTCG | 23 | 47.83 | 61.05 |
| Opisthorsi_10_L | 2 | TCGTCAAGTTAGCTGAACCGC | 21 | 52.38 | 61.03 |
| Opisthorsi_10_R | 2 | CGATAGACAATGGCGCGGTAAT | 22 | 50.00 | 61.36 |
| Opisthorsi_11_L | 1 | GCAGTGATGCGTAAAGTATTGGTG | 24 | 45.83 | 61.04 |
| Opisthorsi_11_R | 1 | TTAAGCGGTCTAGCCACTACCC | 22 | 54.55 | 61.72 |
| Opisthorsi_12_L | 2 | GTTATTACCACTCGTATGACTCCGG | 25 | 48.00 | 61.16 |
| Opisthorsi_12_R | 2 | CAGTCCCGTCAAAGCAGTACAA | 22 | 50.00 | 60.99 |
| Opisthorsi_13_L | 1 | CATCGACTTATGTCTCTCGAAGCT | 24 | 45.83 | 60.58 |
| Opisthorsi_13_R | 1 | TTGACAGGATTCATCAAAAAGGAATGAA | 28 | 32.14 | 60.77 |
| Opisthorsi_14_L | 2 | GAGAGGGAAAGGCAGATGTAGC | 22 | 54.55 | 60.67 |
| Opisthorsi_14_R | 2 | GGTGCAGAAAACAGCTCTCTCT | 22 | 50.00 | 60.73 |
| Opisthorsi_15_L | 1 | TGCCGCAAGATTTTCGAAAGAC | 22 | 45.45 | 60.53 |
| Opisthorsi_15_R | 1 | TGATACAGCCTCGCACTCTTG | 21 | 52.38 | 60.24 |
| Opisthorsi_16_L | 2 | GTTGGATTACTACACCGTCCACC | 23 | 52.17 | 61.18 |
| Opisthorsi_16_R | 2 | ACCGTATTCATTGTCAGGTAGCG | 23 | 47.83 | 60.99 |
| Opisthorsi_17_L | 1 | CTAGAGCAATACAACCGTGCCA | 22 | 50.00 | 60.86 |
| Opisthorsi_17_R | 1 | AGCTAGAAGCACCTCAGTCGTA | 22 | 50.00 | 60.80 |
| Opisthorsi_18_L | 2 | CATGCCTATGGTAGTATGGTACCG | 24 | 50.00 | 60.70 |
| Opisthorsi_18_R | 2 | ACTGTCTGTAGAGCTGAGTACTCC | 24 | 50.00 | 61.19 |
| Opisthorsi_19_L | 1 | CACTGCGCAATCAGTATAGTTTTATCA | 27 | 37.04 | 60.74 |

| | | | | | |
|---|---|---|---|---|---|
| Opisthorsi_19_R | 1 | TCGAGTATACATGTTGATGATACCACA | 27 | 37.04 | 60.48 |
| Opisthorsi_20_L | 2 | CTTGCGTTGATGGTTGAGTGTG | 22 | 50.00 | 60.78 |
| Opisthorsi_20_R | 2 | AACCACCACCGATCATTGACTAC | 23 | 47.83 | 60.87 |
| Opisthorsi_21_L | 1 | CGGCGGTGTACAATTTGCAAG | 21 | 52.38 | 61.09 |
| Opisthorsi_21_R | 1 | AACTCCAAATCAGGCAGCGAC | 21 | 52.38 | 61.32 |
| Opisthorsi_22_L | 2 | TCGCTTCCGAAGAACAGAGAGA | 22 | 50.00 | 61.32 |
| Opisthorsi_22_R | 2 | GGTCGAAAAATAGTTTCTAAAGAACTTCG | 29 | 34.48 | 60.32 |
| Opisthorsi_23_L | 1 | TGAATGGTCCTATGGGCTCAGT | 22 | 50.00 | 61.09 |
| Opisthorsi_23_R | 1 | ACCAGTGCAAACGTACGTACTC | 22 | 50.00 | 61.04 |
| Opisthorsi_24_L | 2 | CCCATTCTTTTTCCTGCGGTGA | 22 | 50.00 | 61.58 |
| Opisthorsi_24_R | 2 | GGGTACTCATAAACAGGAAGATGAAAC | 27 | 40.74 | 60.42 |
| Opisthorsi_25_L | 1 | GTCTTGCGTGATTCTTCTACGGT | 23 | 47.83 | 61.17 |
| Opisthorsi_25_R | 1 | AGCACCTTCACGTCTTTCACC | 21 | 52.38 | 60.91 |
| Opisthorsi_26_L | 2 | CCACTGCTTGAGTTGCTTTTGG | 22 | 50.00 | 60.98 |
| Opisthorsi_26_R | 2 | ATCGGCTATATCTTGAAGCGCG | 22 | 50.00 | 61.16 |
| Opisthorsi_27_L | 1 | AAGGGGCCTGATATTGTGCG | 20 | 55.00 | 60.48 |
| Opisthorsi_27_R | 1 | GCGACGTGAAGTAAGTATGTAATGC | 25 | 44.00 | 60.92 |
| Opisthorsi_28_L | 2 | TTTGGGTGCGATTTTTCTTCTGC | 23 | 43.48 | 61.05 |
| Opisthorsi_28_R | 2 | ACATCGTCATAGAGAGAAAAGTAGTCAG | 28 | 39.29 | 60.96 |
| Opisthorsi_29_L | 1 | TACCGTTATCAACCACCACACC | 22 | 50.00 | 60.47 |
| Opisthorsi_29_R | 1 | AGGGCCAGTTAGATGAGACGAT | 22 | 50.00 | 60.88 |
| Opisthorsi_30_L | 2 | CCTGCGGAATGGAAAGTTGAATTT | 24 | 41.67 | 60.88 |
| Opisthorsi_30_R | 2 | TGAAAATAAAAGTCCGATAGCAGCG | 25 | 40.00 | 60.59 |

**Table S4.**

Primers used for complete genome sequencing of OPLV.

| Name | Pool | Sequence | Size (bp) | %GC | Tm (°C) |
|---|---|---|---|---|---|
| OPLV_seg1_1_L | 1 | TGAGAAATCAGGTTGTATTGTTACATCG | 28 | 35.71 | 60.70 |
| OPLV_seg1_1_R | 1 | GGATGTTAGGAACTATATCAGACATGGC | 28 | 42.86 | 61.80 |
| OPLV_seg1_2_L | 2 | CACCACAAGATGTTCTAAAATTTCCTCA | 28 | 35.71 | 61.02 |
| OPLV_seg1_2_R | 2 | AGGGTTCTTCCTTCTCAATTCATCC | 25 | 44.00 | 60.96 |
| OPLV_seg1_3_L | 1 | GTTCTCAACATTTGCCATTGATGAAG | 26 | 38.46 | 60.34 |
| OPLV_seg1_3_R | 1 | CCATAAGAGGTGGGTATAATTAAAGGAAAC | 30 | 36.67 | 60.88 |
| OPLV_seg1_4_L | 2 | AACAAGCACTCAACAAGCAGAC | 22 | 45.45 | 60.08 |
| OPLV_seg1_4_R | 2 | ACTGCGTTTAAATTTCTTGCCCC | 23 | 43.48 | 60.81 |
| OPLV_seg1_5_L | 1 | CCTGTTGAATTTGTTTCATCCTTGC | 25 | 40.00 | 60.20 |
| OPLV_seg1_5_R | 1 | TGCAGAACAAGCAATTTCTCTAGTC | 25 | 40.00 | 60.20 |
| OPLV_seg1_6_L | 2 | CATTGAGAAATTCATTGAAACTTTCCCAG | 29 | 34.48 | 60.92 |
| OPLV_seg1_6_R | 2 | GACTTAAAGCTGTACATTCTGATGTTGT | 28 | 35.71 | 60.86 |
| OPLV_seg1_7_L | 1 | AAACCCACCAAATCTCATCAGCA | 23 | 43.48 | 61.01 |
| OPLV_seg1_7_R | 1 | TTCTTGACCTGGGACATGATGG | 22 | 50.00 | 60.21 |
| OPLV_seg1_8_L | 2 | GGGATTCTTCTAGCCCTGGACA | 22 | 54.55 | 61.41 |
| OPLV_seg1_8_R | 2 | GTTTGATATTCCAACTCAGGTGCAG | 25 | 44.00 | 60.77 |
| OPLV_seg1_9_L | 1 | AGGAAGTCCTCACATTAATGTATACAGG | 28 | 39.29 | 60.98 |
| OPLV_seg1_9_R | 1 | GCACTCCTTTAAAACAAGCAGAAC | 24 | 41.67 | 59.57 |
| OPLV_seg1_10_L | 2 | GCTAGTAATGTTGATCTTGAGGAAGC | 26 | 42.31 | 60.45 |
| OPLV_seg1_10_R | 2 | TCTGATCTAGGACTTGACCTCCTG | 24 | 50.00 | 60.96 |
| OPLV_seg1_11_L | 1 | ATGACCCACCCAGAAGTCAAAC | 22 | 50.00 | 60.67 |
| OPLV_seg1_11_R | 1 | ATCCTTGCATCATGCCTGAGTC | 22 | 50.00 | 60.93 |
| OPLV_seg1_12_L | 2 | TGGATGCTCAAGAAATAGTTCATGG | 25 | 40.00 | 59.79 |
| OPLV_seg1_12_R | 2 | TCTTTACTGCTGATGATCCCAAGC | 24 | 45.83 | 61.25 |
| OPLV_seg1_13_L | 1 | TCATATCAAATGCCCCTATACAATACGT | 28 | 35.71 | 61.08 |
| OPLV_seg1_13_R | 1 | TGGGGTCAGGAACTCTTAGAAGT | 23 | 47.83 | 60.70 |
| OPLV_seg1_14_L | 2 | GGTTTGTATTTCTATAGAATGGCAGGC | 27 | 40.74 | 61.01 |
| OPLV_seg1_14_R | 2 | TTCATGGCGATGTGAACCTCA | 21 | 47.62 | 60.10 |
| OPLV_seg1_15_L | 1 | ACAAATGTCAGAAAATACAAGGAACTCA | 28 | 32.14 | 60.50 |
| OPLV_seg1_15_R | 1 | ACTGCTCATGAAGAGGGAAAAACA | 24 | 41.67 | 61.01 |
| OPLV_seg1_16_L | 2 | GGATGAGCAAGAGCCCTTTATCA | 23 | 47.83 | 60.69 |
| OPLV_seg1_16_R | 2 | TTTCCCCTTCTTGGTGGTCCTA | 22 | 50.00 | 60.89 |
| OPLV_seg1_17_L | 1 | CCCTTTCTGAGAGACACGTTCTAC | 24 | 50.00 | 60.93 |
| OPLV_seg1_17_R | 1 | ACCTCTTGTCACAAGAGCTTTGT | 23 | 43.48 | 60.63 |

| | | | | | |
|---|---|---|---|---|---|
| OPLV_seg1_18_L | 2 | CTTTTCTGACAAGCGCAAGGC | 21 | 52.38 | 61.03 |
| OPLV_seg1_18_R | 2 | TGGATTCTATAACTGGTTCCACTTTCC | 27 | 40.74 | 61.30 |
| OPLV_seg1_19_L | 1 | CAGACAACTTAGAAGTGTTGCAGC | 24 | 45.83 | 60.87 |
| OPLV_seg1_19_R | 1 | GCTCAGGCAAGTTCTCTGACAT | 22 | 50.00 | 60.80 |
| OPLV_seg1_20_L | 2 | AGACTTGGTTGCAGAGTGTGAAA | 23 | 43.48 | 60.88 |
| OPLV_seg1_20_R | 2 | GAGGAGTTGACAGTTGGAAAGCA | 23 | 47.83 | 61.25 |
| OPLV_seg1_21_L | 1 | GCACTGATGGAACCACTAGATCC | 23 | 52.17 | 61.00 |
| OPLV_seg1_21_R | 1 | ACTTCTTTTCTTCAGACGAGGTGG | 24 | 45.83 | 61.06 |
| OPSV_seg2_1_L | 1 | GGACACACAGACACCCATACAAA | 23 | 47.83 | 61.00 |
| OPSV_seg2_1_R | 1 | TGTTCTCAGGCGTGACAGTCTA | 22 | 50.00 | 61.26 |
| OPSV_seg2_2_L | 2 | ATGTCTGTGGTCTTCCTCCTGA | 22 | 50.00 | 60.68 |
| OPSV_seg2_2_R | 2 | ATCCTGGGCTGCTTTCACAAAT | 22 | 45.45 | 61.01 |
| OPSV_seg2_3_L | 1 | TGTATGCAGTTGCCTTGGGTG | 21 | 52.38 | 61.26 |
| OPSV_seg2_3_R | 1 | TCTGGTCCTACTCTTCACTCTGG | 23 | 52.17 | 60.82 |

**Table S5.**

Detailed descriptive information of the viruses discovered in *O. viverrini* and *S. haematobium*

| Virus Name | Genbank Accession | Abreviation | Host | Length (nt) | Best blastx hit | Acc | % id | Phylum | Class | Order | Family | Genus |
|---|---|---|---|---|---|---|---|---|---|---|---|---|
| **Schistohaemavirus** | BK075116 | ShV | *Schistosoma haematobium* | 9179 | Dicrocoelium Nege-like virius | WFD53207.1 | 53,8 | *Kitrinoviricota* | *Alsuviricetes* | *Martellivirales* | "Prenviridae" | |
| **Schistohaema togalivirus** | BK075117 | ShTLV | *Schistosoma haematobium* | 14596 | Dicrocoelium unclassified virus 2 | WFD53224.1 | 38,8 | *Kitrinoviricota* | *Alsuviricetes* | *Martellivirales* | "Togaliviridae" | |
| **Schistohaemarhabdovirus** | BK075118 | ShRV | *Schistosoma haematobium* | 13053 | Alphaplatrhavirus turkestanicum | DAZ87982.1 | 89,6 | *Negarnaviricota* | *Monjiviricetes* | *Mononegavirales* | *Rhabdoviridae* | *Alphaplatrhavirus* |
| **Schistohaemendornavirus** | BK075119 | ShMV | *Schistosoma haematobium* | 13705 | Schistomendorna virus | DAZ87976.1 | 40 | *Kitrinoviricota* | *Alsuviricetes* | *Martellivirales* | *Endornaviridae* | *Alphaendornavirus* |
| **Opisthorsivirus** | PX443621 | OSV | *Opisthorchis viverrini* | 9548 | Clonorsi virus 3 | DAZ87859.1 | 91 | *Kitrinoviricota* | *Alsuviricetes* | *Martellivirales* | "Prenviridae" | |
| **Opisthorhabdovirus** | PX443622 | ORV | *Opisthorchis viverrini* | 10986 | Clonorhabdovirus 2 | DAZ87848.1 | 97,3 | *Negarnaviricota* | *Monjiviricetes* | *Mononegavirales* | *Rhabdoviridae* | *Gammaplatrhavirus* |
| **Opisthorphenuili virus** | PX443623 | OPLV | *Opisthorchis viverrini* | RNA1: 6349 | Clonorchi Phenuili virus | DAZ87849.1 | 89,1 | *Negarnaviricota* | *Bunyaviricetes* | *Hareavirales* | "Phenuiliviridae" | |
| | | | | RNA2: 1076 | Clonorchi Phenuili virus | DAZ87850.1 | 87,1 | *Negarnaviricota* | *Bunyaviricetes* | *Hareavirales* | "Phenuiliviridae" | |

**Table S6**

Primer sequences used for Rt-qPCR detection of *O. viverrini* viruses.

| Primer name | Sequence | System and amplicon length |
|---|---|---|
| ParaRhabdo-N_F | GCCATTCAGTCYGTGTGGATG | qPCR pan-rhabdo fluke-borne viruses – Target : nucleoprotein |
| ParaRhabdo-N_R | TTRCAGAGTGCGTCGATAGTCG | Tm 60°C - 146 bp |
| ParaRhabdo-G_F | TGAAGCTGTGACCATCTGGATG | qPCR pan-rhabdo fluke-borne viruses – target: glycoprotein |
| ParaRhabdo-G_R | TYTTAGCAAATCTAAGACATATTGCAG | Tm 60°C - 187 bp |
| ParaRhabdo-L_F | CATCATCRTGGTATAAGTGGATGG | qPCR pan-rhabdo fluke-borne viruses – target: RNA dependent RNA polymerase |
| ParaRhabdo-L_R | ACYCTACGTTTACTTGGGTCTG | Tm 60°C - 142 bp |
| ParaBunya-L_F | GTYATAAAYACTCATGCAGTCACATC | qPCR pan-bunya fluke-borne viruses – target : RNA dependent RNA polymerase |
| ParaBunya-L_R | CTAACWATCTCCTTTCTATCCATCCA | Tm 60°C - 230 bp |
| ParaBunya-S_F | CGTGATTACATCACAAATGAATAATCC | qPCR pan-bunya fluke-borne viruses – target: nucleoprotein |
| ParaBunya-S_R | CAAARCCTTCRTATCTGTACAGGTC | Tm 60°C - 212 bp |
| ParaMartelli-M_L | GTKCTTGTRCTYGTGGCCG | qPCR pan-martelli fluke-borne viruses - Tm 58°C - 152 bp |
| ParaMartelli-M_R | GGRRYRAAHAYAACCATACCTGC | |
| ParaMartelli-L_F2 | CATGAGMGTGARGTCTKTKGGG | qPCR pan-martelli fluke-borne viruses - Tm 60°C - 111 bp |
| ParaMartelli-L_R | CTCGWCCCGCATAGATGGTG | |
| OvActin_F | GGCAGATTCCATACCCAAGA | qPCR *O. viverrini* Actin target : Actin |
| OvActin_R | CGAGCGTGGTTACAGTTTCA | Tm 55 °C – 230 bp |

**Table S7**
Primers used to generate synthetic RNA for viral RNA quantification of fluke viruses

| OligoName | Sequence | Tm - amplicon length |
|---|---|---|
| Oviv-Rhabdo-L_T7-F | TAATACGACTCACTATAGGGACAGTCCACGACCACTC | Tm 58°C - 229 bp +18 bp T7 promotor |
| Oviv-Rhabdo-L_T7-R | TCTGTATCGACTCCATATGGTC | |
| Oviv-Bunya-S_T7-F | TAATACGACTCACTATAGGGACACACAGACACCCATAC | Tm 58°C - 279 bp +18 bp T7 promotor |
| Oviv-Bunya-S_T7-R | TTCTAGCCTTCATCACCAAGTAC | |
| Oviv-Martelli-L_T7-F | TAATACGACTCACTATAGGGAAACACGATCATACAATGTC | Tm 58°C - 193 pb (+17 pb T7 promotor) |
| Oviv-Martelli-L_T7-R | GATCCATCAACAACTTCGAAACC | |

**Table S8.**

RNAscope probes position ordered to biotechne ®

| NPR | Catalog # | Probe Name | Target Region |
|---|---|---|---|
| NPR-0053382 | 157303* | V-ORV-RdRP-O1 | 809-1727 |
| NPR-0053383 | 157302* | V-OSV-ORF1-O1 | 1198-2107 |
| NPR-0053384 | 157301* | V-OPLV-L-O1 | 620-5466 |
| NPR-0058503 | 180618* | V-ORV-O1-sense | reverse-complement sequence of 28-987 |
| NPR-0058505 | 180620* | V-OSV-O1-sense | reverse-complement sequence of 902-1865 |
| NPR-0058506 | 180621* | V-OPLV-O1-sense | reverse-complement sequence of 2-1894 |

**Table S9.**

Two-sided Wilcoxon-Mann-Whitney tests for differences in IgG titers against OPLV, ORV, OSV and *Ov*-TSP-2 between French non-endemic subjects (FR, n=100) and Thai *O. viverrini* infected subjects (Ov-Thai, n=455). Reported effect size was calculated as r=Z/sqrt(n), where n is the total number of observations; p-values were adjusted using Benjamini-Hochberg correction. CI: confidence interval; IQR: interquartile range

| Titer | **Group 1** median (IQR) | **Group 2** median (IQR) | Z statistic | p-value | Adjusted p-value | **Effect size** (95%CI) | Magnitude |
|---|---|---|---|---|---|---|---|
| **OPLV** | FR<br>4.9 (1.9) | Ov-Thai<br>7.7 (3.2) | 8027.5 | 3.69e-24 | **3.69e-24** **** | 0.43 (0.37-0.49) | moderate |
| **OSV** | FR<br>5.0 (1.4) | Ov-Thai<br>7.5 (3.3) | 6112.0 | 2.13e-30 | **8.52e-30** **** | 0.49 (0.43-0.54) | moderate |
| **ORV** | FR<br>2.8 (1.5) | Ov-Thai<br>5.8 (3.5) | 7789.5 | 6.79e-25 | **9.05e-25** **** | 0.44 (0.38-0.49) | moderate |
| **TSP2** | FR<br>3.1 (1.4) | Ov-Thai<br>6.2 (3.8) | 6436.5 | 2.73e-29 | **5.46e-29** **** | 0.48 (0.42-0.53) | moderate |

**Table S10.**

Receiver-operator characteristics (ROC) curve analysis from the quantification of IgG titers against OPLV, ORV, OSV and *Ov*-TSP-2 in French non-endemic subjects (FR, n=100) and Thai *O. viverrini* infected subjects (Ov-Thai, n=455). The 95% confidence interval, computed by bootstrap using pROC package, is indicated in brackets.

| Titer | Threshold | Accuracy | Sensitivity | Specificity | Positive predicted value | Negative predicted value |
|---|---|---|---|---|---|---|
| **TSP-2** | 4.64 (3.93-5.21) | 0.77 (0.7-0.84) | 0.74 (0.64-0.84) | 0.94 (0.84-1) | 0.98 (0.96-1) | 0.44 (0.37-0.53) |
| **ORV** | 3.94 (3.52-5.08) | 0.75 (0.64-0.82) | 0.74 (0.58-0.82) | 0.85 (0.73-0.97) | 0.96 (0.93-0.99) | 0.41 (0.33-0.5) |
| **OSV** | 6.36 (5.7-6.71) | 0.75 (0.69-0.82) | 0.71 (0.63-0.82 | 0.92 (0.8-0.98) | 0.98 (0.95-0.99) | 0.41 (0.36-0.51) |
| **OPLV** | 6.24 (5.66-6.71) | 0.77 (0.71-0.82) | 0.76 (0.67-0.83) | 0.82 (0.72-0.9) | 0.95 (0.93-0.97) | 0.43 (0.37-0.51) |

**Table S11.**
Kruskal-Wallis tests for differences in IgG titers against OPLV, ORV, OSV and *Ov*-TSP-2 between diagnostic groups: non-endemic French subjects (FR, n=100), Thai subjects without periductal fibrosis (APF-, n=184), with periductal fibrosis (APF+, n=184) and with cholangiocarcinoma (CCA, n=87). Reported effect size was calculated as eta2[H] = (H - k + 1)/(n - k); p-values were adjusted using Benjamini-Hochberg correction. CI: confidence interval; IQR: interquartile range; df: degrees of freedom

| **Titer** | **Groups compared** median (IQR) | **H statistic** | **df** | **p-value** | **Adjusted p-value** | **Effect size** (95%CI) | **Magnitude** |
|---|---|---|---|---|---|---|---|
| **OSV** | APF- 6.9 (3.0) | 156.4 | 3 | 1.09e-33 | **1.45e-33** **** | 0.28 (0.22-0.34) | large |
| | APF+ 8.2 (4.2) | | | | | | |
| | CCA 7.4 (2.6) | | | | | | |
| | FR 5.0 (1.4) | | | | | | |
| **ORV** | APF- 4.7 (3.5) | 200.7 | 3 | 3.05e-43 | **1.22e-42** **** | 0.36 (0.3-0.42) | large |
| | APF+ 7.2 (3.0) | | | | | | |
| | CCA 5.2 (2.9 | | | | | | |
| | FR 2.8 (1.5) | | | | | | |
| **TSP2** | APF- 6.2 (4.3) | 127.1 | 3 | 2.33e-27 | **2.33e-27** **** | 0.23 (0.18-0.28) | large |
| | APF+ 6.2 (3.2) | | | | | | |
| | CCA 6.5 (4.3) | | | | | | |
| | FR 3.1 (1.4) | | | | | | |
| **OPLV** | APF- 7.0 (3.1) | 163.1 | 3 | 3.98e-35 | **7.96e-35** **** | 0.29 (0.23-0.36 | large |
| | APF+ 8.8 (3.2) | | | | | | |
| | CCA 7.3 (3.3) | | | | | | |
| | FR 4.9 (1.9) | | | | | | |

**Table S12.**
Two-sided Wilcoxon-Mann-Whitney tests for differences in IgG titers against OPLV, ORV, OSV and *v*-TSP-2 between diagnostic groups: non-endemic French subjects (FR, n=100) and Thai subjects without periductal fibrosis (APF-, n=184), with periductal fibrosis (APF+, n=184) and with cholangiocarcinoma (CCA, n=87). Reported effect size was calculated as r=Z/sqrt(n), where n is the total number of observations; p-values were adjusted using Benjamini-Hochberg correction. CI: confidence interval

| **Titer** | **Group 1** | **Group 2** | **Z statistic** | **p-value** | **Adjusted p-value** | **Effect size** (95%CI) | **Magnitude** |
|---|---|---|---|---|---|---|---|
| OSV | APF- | APF+ | 11488.0 | 9.76e-08 | **1.46e-07** **** | 0.28 (0.18-0.37) | small |
| OSV | APF- | CCA | 6875.5 | 0.061 | 0.061 ns. | 0.11 (0.0065-0.23) | small |
| OSV | APF- | FR | 14947.0 | 3.53e-18 | **7.06e-18** **** | 0.52 (0.43-0.59) | large |
| OSV | APF+ | CCA | 9616.0 | 0.007 | **0.009** ** | 0.16 (0.05-0.28) | small |
| OSV | APF+ | FR | 16723.0 | 5.29e-30 | **3.17e-29** **** | 0.68 (0.61-0.73) | large |
| OSV | CCA | FR | 7718.0 | 7.41e-20 | **2.22e-19** **** | 0.67 (0.58-0.75) | large |
| ORV | APF- | APF+ | 7048.0 | 3.56e-22 | **1.07e-21** **** | 0.5 (0.42-0.58) | large |
| ORV | APF- | CCA | 6432.5 | 0.009 | **0.009** ** | 0.16 (0.04-0.27) | small |
| ORV | APF- | FR | 12949.5 | 1.41e-08 | **1.69e-08** **** | 0.34 (0.24-0.43) | moderate |
| ORV | APF+ | CCA | 11709.5 | 7.71e-10 | **1.16e-09** **** | 0.37 (0.27-0.48) | moderate |
| ORV | APF+ | FR | 17449.5 | 9.77e-36 | **5.86e-35** **** | 0.74 (0.69-0.78) | large |
| ORV | CCA | FR | 7311.5 | 1.05e-15 | **2.10e-15** **** | 0.59 (0.48-0.68) | large |
| TSP-2 | APF- | APF+ | 16541.0 | 0.705 | 0.737 ns. | 0.02 (0.0015-0.13) | small |
| TSP-2 | APF- | CCA | 7728.0 | 0.647 | 0.737 ns. | 0.03 (0.0019-0.15) | small |
| TSP-2 | APF- | FR | 14958.0 | 3.04e-18 | **9.12e-18** **** | 0.52 (0.43-0.61) | large |
| TSP-2 | APF+ | CCA | 7801.5 | 0.737 | 0.737 ns. | 0.02 (0.0021-0.16) | small |
| TSP-2 | APF+ | FR | 16759.0 | 2.83e-30 | **1.70e-29** **** | 0.68 (0.61-0.73) | large |
| TSP-2 | CCA | FR | 7346.5 | 4.85e-16 | **9.70e-16** **** | 0.59 (0.47-0.7) | large |
| OPLV | APF- | APF+ | 8662.0 | 5.46e-16 | **1.64e-15** **** | 0.42 (0.33-0.51) | moderate |
| OPLV | APF- | CCA | 6429.0 | 0.009 | **0.009** ** | 0.16 (0.05-0.27) | small |
| OPLV | APF- | FR | 13434.5 | 1.50e-10 | **2.25e-10** **** | 0.38 (0.27-0.47) | moderate |
| OPLV | APF+ | CCA | 10397.0 | 7.13e-05 | **8.56e-05** **** | 0.24 (0.11-0.35) | small |
| OPLV | APF+ | FR | 16759.0 | 2.83e-30 | **1.70e-29** **** | 0.68 (0.61-0.74) | large |
| OPLV | CCA | FR | 7279.0 | 2.15e-15 | **4.30e-15** **** | 0.58 (0.47-0.68) | large |

**Table S13.**
Kruskal-Wallis tests for differences in IgG avidity against OPLV, ORV, OSV and *Ov*-TSP-2 between parasitized groups: Thai subjects without periductal fibrosis (APF-, n=72), with periductal fibrosis (APF+, n=72) and with cholangiocarcinoma (CCA, n=71). Reported effect size was calculated as eta2[H] = (H - k + 1)/(n - k); p-values were adjusted using Benjamini-Hochberg correction. CI: confidence interval; IQR: interquartile range; df: degrees of freedom

| **Titer** | **Groups compared** median (IQR) | **H statistic** | **df** | **p-value** | **Adjusted p-value** | **Effect size** (95%CI) | **Magnitude** |
|---|---|---|---|---|---|---|---|
| **OSV avidity** | APF- 43.1 (15.1) | 115.5 | 2 | 8.12e-26 | **1.08e-25** **** | 0.54 (0.43-0.63) | large |
| | APF+ 55.8 (17.6) | | | | | | |
| | CCA 74.2 (6.5) | | | | | | |
| **ORV avidity** | APF- 43.2 (15.2) | 127.1 | 2 | 2.51e-28 | **5.02e-28** **** | 0.59 (0.5-0.67) | large |
| | APF+ 51.2 (16.6) | | | | | | |
| | CCA 75.6 (9.3) | | | | | | |
| **TSP-2 avidity** | APF- 43.7 (17.6) | 89.4 | 2 | 3.83e-20 | **3.83e-20** **** | 0.41 (0.31-0.51) | large |
| | APF+ 40.9 (13.0) | | | | | | |
| | CCA 59.7 (8.4) | | | | | | |
| **OPLV avidity** | APF- 43.0 (14.2 | 133.4 | 2 | 1.09e-29 | **4.36e-29** **** | 0.62 (0.55-0.68) | large |
| | APF+ 57.8 (21.9) | | | | | | |
| | CCA 75.9 (8.3) | | | | | | |

**Table S14.**

Two-sided Wilcoxon-Mann-Whitney tests for differences in IgG avidity against OPLV, ORV, OSV and *v*-TSP-2 between parasitized groups: Thai subjects without periductal fibrosis (APF-, n=72), with periductal fibrosis (APF+, n=72) and with cholangiocarcinoma (CCA, n=71). Reported effect size was calculated as r=Z/sqrt(n), where n is the total number of observations; p-values were adjusted using Benjamini-Hochberg correction. CI: confidence interval

| Titer | Group 1 | Group 2 | Z statistic | p-value | Adjusted p-value | Effect size (95%CI) | Magnitude |
|---|---|---|---|---|---|---|---|
| **OSV av.** | APF- | APF+ | 1413 | 2.49e-06 | **2.49e-06** **** | 0.39 (0.26-0.53) | moderate |
| **OSV av.** | APF- | CCA | 211 | 2.91e-21 | **8.73e-21** **** | 0.79 (0.7-0.85) | large |
| **OSV av.** | APF+ | CCA | 560 | 7.83e-16 | **1.17e-15** **** | 0.67 (0.56-0.76) | large |
| **ORV av.** | APF- | APF+ | 1611 | 8.94e-05 | **8.94e-05** **** | 0.33 (0.18-0.48) | moderate |
| **ORV av.** | APF- | CCA | 105 | 4.42e-23 | **1.33e-22** **** | 0.83 (0.77-0.86) | large |
| **ORV av.** | APF+ | CCA | 338 | 3.45e-19 | **5.17e-19** **** | 0.75 (0.66-0.81) | large |
| **TSP-2 av.** | APF- | APF+ | 2847 | 0.309 | 0.309 ns. | 0.08 (0.0048-0.23) | small |
| **TSP-2 av.** | APF- | CCA | 771 | 5.81e-13 | **8.72e-13** **** | 0.6 (0.47-0.71) | large |
| **TSP-2 av.** | APF+ | CCA | 322 | 1.92e-19 | **5.76e-19** **** | 0.75 (0.67-0.82) | large |
| **OPLV av.** | APF- | APF+ | 1356 | 7.95e-07 | **7.95e-07** **** | 0.41 (0.28-0.54) | moderate |
| **OPLV av.** | APF- | CCA | 27 | 1.81e-24 | **5.43e-24** **** | 0.85 (0.83-0.86) | large |
| **OPLV av.** | APF+ | CCA | 410 | 4.62e-18 | **6.93e-18** **** | 0.72 (0.63-0.8) | large |

**Table S15.**

Random forest models performances: predicting control vs parasitized with or without TSP2 IgG titer. Reported performances were evaluated on independent validation group (FR=30, Ov-Thai=140). The 95% confidence interval computed by Clopper-Pearson exact binomial method via caret package is indicated in brackets.

| | **fluke viral antigens only** | **fluke viral antigens + TSP2** |
|---|---|---|
| **Accuracy** | 0.89 (0.84, 0.94) | 0.94 (0.89, 0.97) |
| **Sensitivity** | 0.63 (0.44, 0.80) | 0.83 (0.65, 0.94) |
| **Specificity** | 0.95 (0.90, 0.98) | 0.96 (0.92, 0.99) |
| **Positive predictive value** | 0.73 (0.52, 0.88) | 0.83 (0.64, 0.94) |
| **Negative predicted value** | 0.92 (0.87, 0.96) | 0.96 (0.92, 0.99) |

**Table S16.**
Random forest models performances predicting parasitized groups diagnostics: Thai subjects without periductal fibrosis (APF-, n=72), with periductal fibrosis (APF+, n=72) and with cholangiocarcinoma (CCA, n=71). We used a random forest model to classify subjects based on IgG titers and IgG avidity against OPLV, ORV, OSV and *v*-TSP-2, using a training set of **147** subjects (APF-=49, APF+=49, CCA=49) and a validation set of **68** subjects (APF-=23, APF+=23, CCA=22). The 95% confidence interval is indicated in brackets.

| | **CCA** | **APF-** | **AFP+** |
|---|---|---|---|
| **Accuracy** | 0.94 (0.86, 0.98) | 0.94 (0.86, 0.98) | 0.90 (0.80, 0.96) |
| **Sensitivity** | 1.00 (0.85, 1.00) | 0.91 (0.72, 0.99) | 0.78 (0.56, 0.93) |
| **Specificity** | 0.91 (0.79, 0.98) | 0.98 (0.88, 1.00) | 0.96 (0.85, 0.99) |
| **Positive predictive value** | 0.85 (0.65, 0.96) | 0.95 (0.77, 1.00) | 0.90 (0.68, 0.99) |
| **Negative predicted value** | 1.00 (0.92, 1.00) | 0.96 (0.85, 0.99) | 0.90 (0.77, 0.97) |

**Table S17.**
Feature importance for all Random forest models. The mean decrease Gini is expressed as a percentage

| Features | Viral titers only | Viral titers + TSP2 | FR vs APF- vs APF+ | CCA vs APF- vs APF+ |
|---|---|---|---|---|
| TSP2 | - | **41.92** | 40.67 | 6.95 |
| ORV | 30.94 | 22.11 | **46.80** | **14.10** |
| OSV | 26.88 | 20.53 | 34.94 | 8.32 |
| OPLV | **37.03** | 26.07 | 34.87 | 4.44 |
| TSP2 (avidity) | - | - | - | 10.70 |
| ORV (avidity | - | - | - | **12.74** |
| OSV (avidity) | - | - | - | 11.97 |
| OPLV (avidity) | - | - | - | 12.16 |
